# Different dopaminergic neurons signal absolute and relative aversive value in the *Drosophila* mushroom body

**DOI:** 10.1101/2022.02.02.478814

**Authors:** Maria E. Villar, Miguel Pavão-Delgado, Marie Amigo, Pedro F. Jacob, Nesrine Merabet, Anthony Pinot, Sophie A. Perry, Scott Waddell, Emmanuel Perisse

## Abstract

Animals use prior experience to assign absolute (good or bad) and also relative (better or worse) value to new experience. These learned values guide appropriate later decision-making. While our understanding of how the dopaminergic system computes absolute value is relatively advanced, the mechanistic underpinnings of relative valuation are unclear. Here we reveal mechanisms of absolute and relative aversive valuation in *Drosophila*. Three types of punishment-sensitive dopaminergic neurons (DANs) drive intensity-scaled plasticity at their respective mushroom body output neuron (MBON) connections to code absolute aversive value. In contrast, by comparing current and previous aversive experiences the MBON-DAN network can code relative aversive value by recruiting a specific subtype of reward-coding dopaminergic neurons which assigns a ‘better than’ value to the lesser of two aversive experiences. This study therefore provides an important functional consequence of having opposing populations of DANs and illustrates how these can operate together as a system within the MB network to code and compare sequential aversive experience to learn relative aversive value.

## Introduction

Value-based decisions require animals to make choices between several options based on a prediction of their relative subjective value learned through prior experience of these options^1^. Associative learning provides a means to assign absolute (good or bad) values to experience that can be used to guide appropriate approach or avoidance behavior at a later time ^2^. During learning, animals can also compare the value of their current experience with that of prior knowledge and assign a relative value (better or worse) between these experiences to promote a more accurate future economic-based choice ^3–5^. While a substantial body of research has investigated mechanisms for relative reward-value coding ^3,6–15^, we know less about how relative aversive value is computed during learning to guide appropriate value-based decisions ^5,16–19^. Reinforcement learning models propose that learning occurs when actual outcome value differs from predicted value ^20,21^. This process and the error computed between actual and predicted value is driven by a valuation circuit that includes dopaminergic and GABAergic neurons ^22–24^. However, it is unclear whether similar circuits can compare current and previous experience to assign relative aversive value to sensory stimuli during learning.

Anatomically discrete dopaminergic neurons (DANs) in *Drosophila* and mice provide either positive or negative teaching signals during learning (for reviews see ^25–28)^. In flies, these different DANs project to unique compartments of the mushroom body (MB), a central brain structure essential for olfactory learning and memory as well as several goal-directed behaviors ^29–32^. DANs from the protocerebral posterior lateral 1 (PPL1) cluster mostly projecting to the vertical lobes of the MB relay punishment and signal negative value to experience during learning ^33–38^. Many DANs from the protocerebral anterior medial (PAM) cluster projecting to the horizontal lobes of the MB assign positive reward value to experience during learning ^39–46^. Sparse activation of subpopulations of the MB intrinsic cells, the Kenyon cells (KCs), receiving odorant stimulation perceived in the antennae and maxillary palps, allow a specific representation of olfactory memories ^17,47,48^. MB KCs synapse onto mushroom body output neurons (MBONs), which project into downstream structures ^49^ to drive (for most of them) approach or avoidance behavior ^32,44,50^ but see ^51^. Some MBONs are synaptically interconnected, providing cross excitation or inhibition between MB compartments. Lastly, many MBONs make synapses outside the MB onto DAN dendrites in a feedback or feedforward manner ^31,44,46,52–55^. During olfactory associative learning, specific DANs provide punishment or reward signals to individual MB compartments, and the released dopamine depresses synaptic strengths between sparse odor-activated KCs and the MBONs whose dendrites reside within the relevant compartments ^50,52,56–63^. As a result, learning-induced plasticity within different MB compartments reconfigures the MBON ensemble output signal to promote either learned approach or avoidance behavior ^32,64^. Olfactory aversive learning reduces the odor drive of approach-directing MBONs, hence tilting the MBON network towards promoting odor avoidance. In contrast, appetitive learning reduces the odor drive of avoidance-directing MBONs, leaving the network in a configuration that promotes odor approach.

Flies have the capacity to perceive, learn and compare differences in the intensity of punishment and adapt their behavior accordingly ^17,65–69^. Here, we combined genetic interventions with behavioral analyses, anatomical characterization, and *in vivo* 2-photon calcium imaging to investigate the detailed circuit requirements that allow flies to write and compare olfactory aversive memories of different intensities during learning to promote appropriate value-based choices. We found that three types of aversive PPL1 DANs, PPL1-γ1ped, PPL1-γ2α’1 and PPL1-α2α’2 show differential responses to electric shock punishment of varying intensity. As a result, the intensity of shock reinforcement correlates with the magnitude of learning-driven plasticity at the corresponding KC to MBON-γ1ped>αβ, MBON-γ2α’1 and MBON-α2sc junctions. We next used a specific behavioral paradigm in which flies associate three odors with 0V, 60V and 30V punishment respectively to identify the circuits involved in coding relative aversive value. Loss of function screening revealed a role for the aversive PPL1-γ1ped, PPL1-γ2α’1 and PPL1-α2α’2 DANs in addition to the rewarding PAM-β’2aγ5n DANs during learning of relative aversive value. Strikingly, the PAM-β’2aγ5n DANs were only required during the last odor-punishment association when comparison can be made with the previous odor-punishment association. Recording from PAM-β’2aγ5n DANs during learning revealed that these neurons signal relative aversive value by increasing their responsiveness when the odor-low shock association is better than a previous odor-high shock association. We also recorded from MBON-γ1ped>αβ, MBON-γ2α’1 and MBON-γ5β’2a which make synaptic connections onto PAM-β’2aγ5n DANs. This revealed a positive difference in the odor-responses of MBON-γ2α’1 between the current and previous aversive experience. The increased responsiveness of cholinergic MBON-γ2α’1 is likely to provide the excitatory input that is necessary to drive the PAM-β’2aγ5n DANs ‘better than’ value signal for the less aversive experience, and thereby learning the relative aversive value.

## Results

### Individual PPL1 DANs respond differently to electric shock intensity

Flies can learn to associate odors with different intensities of electric shocks, reaching a plateau of memory performance around 60V (**Fig. 1a-b**; ^65–67,70^). However, it is currently unknown how shock-responsive DANs in the PPL1 cluster assign aversive values of different intensities to odors during learning. It has been previously suggested that electric shock intensity (i.e. different current intensity) is represented in a graded fashion in PPL1 DANs ^71,72^. To verify these findings in a setting resembling standard training conditions (i.e. with different voltage settings), we used GAL4 ^73^ driven expression of UAS-GCaMP6m ^74^ to measure calcium responses in punishment DANs following electric shocks of 0V, 15V, 30V, 45V, 60V or 90V delivered to the fly legs (**Fig. 1c**). We focused on three DANs from the PPL1 cluster, two of which - PPL1-γ1ped (c061;MBGAL80-GAL4) and PPL1-γ2α’1 (MB296B-GAL4)- are known to assign negative value to odors during learning ^34,36^ while the third - PPL1-α2α’2 (MB058B-GAL4) - does not exhibit clear immediate reinforcing properties when artificially triggered alone ^37,58^. We used 2-photon microscopy to monitor DAN responses in individual flies while they received 12 presentations of 1.5 s electric shocks at 0.2Hz on the legs, a protocol commonly used for aversive conditioning (**Fig. 1a-c**; ^65^). We found that calcium transients in both PPL1-γ1ped and PPL1-γ2α’1, but not PPL1-α2α’2 DANs, significantly increased with electric shock stimulation above 30V, fitting with a linear regression (magenta) and correlating with shock intensity (r value in black) (**Fig. 1d-i**). Although not reaching significance, PPL1-α2α’2 DAN responses seem to be synchronized with shock stimulations of 60V and 90V, hence a fitted regression line with a slope different from 0 (magenta). For all DANs, the increased response was observable from the first of the twelve shock presentations for each intensity (**Supplementary Fig. 1a-f**). While proportional shock responses were readily apparent in PPL1-γ1ped DANs (**Fig. 1d-e**), PPL1-γ2α’1 DAN responses remained stable from 30 to 90V (**Fig. 1f-g**). Accordingly, when we compared the slopes of the corresponding fitted regression lines for each DAN response (in magenta in **Fig. 1e, g and i**), we found a significant difference between PPL1-γ1ped and PPL1-γ2α’1, and between PPL1-γ1ped and PPL1-α2α’2 DAN responses, whereas PPL1-γ2α’1 and PPL1-α2α’2 DANs were indistinguishable (**Supplementary Fig. 1g**). We noted that PPL1-γ1ped and PPL1-γ2α’1 DANs respond similarly during the first shock (**Supplementary Fig. 1h**). Overall, these data show that PPL1-γ1ped, PPL1-γ2α’1 and PPL1-α2α’2 DANs respond differently to a range of electric shock intensities and demonstrate a functional segregation between these three types of PPL1 DANs.

**Fig. 1.**
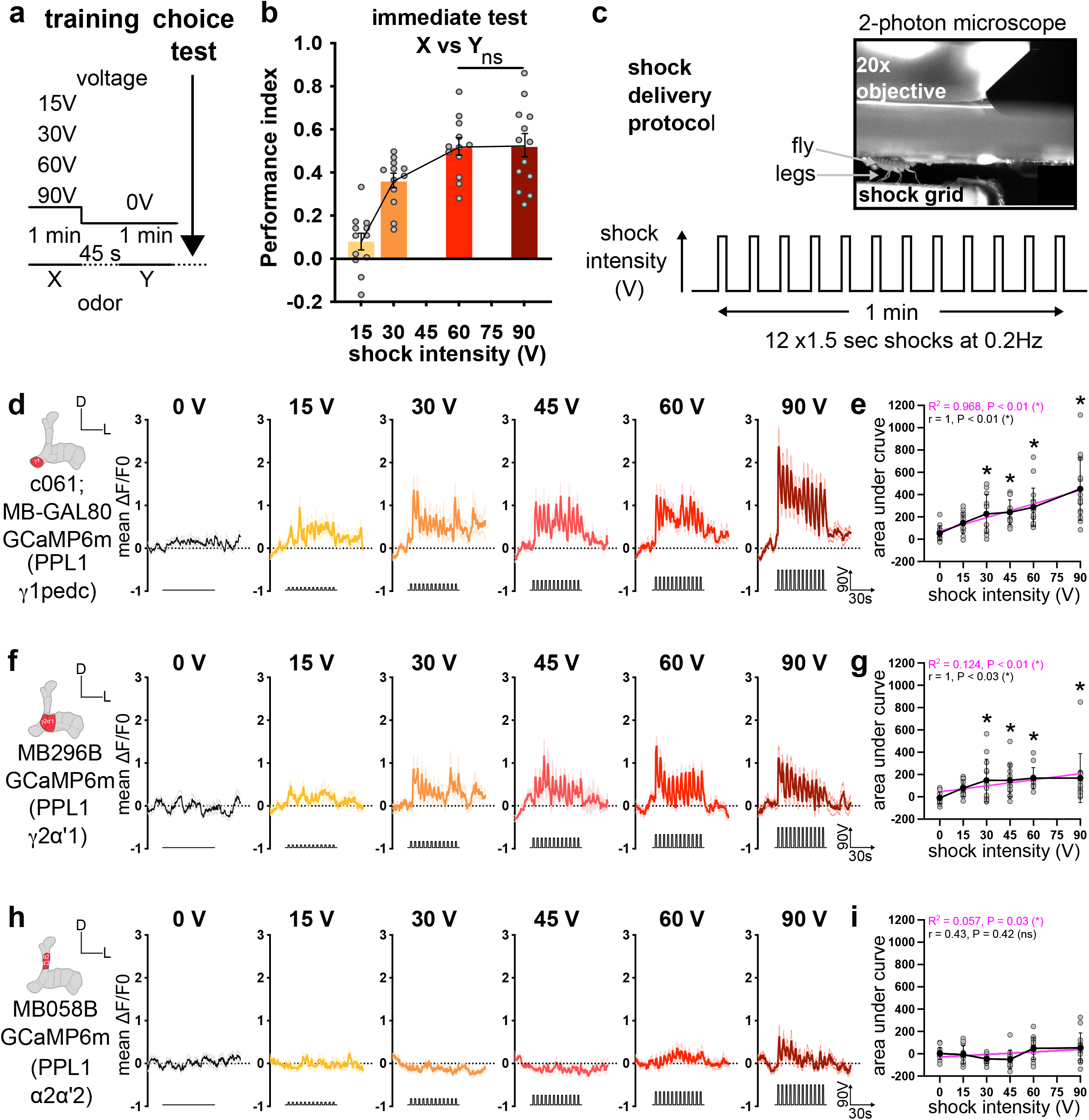
PPL1-γ1ped, PPL1-γ2α’1 and PPL1-α2α’2 DANs respond differently to electric shock intensity. **a.** Training procedure of differential conditioning; Odor X is paired with either 15V, 30V, 60V or 90V. Odor Y is presented alone. **b.** Memory performance immediately after training. Flies choose between odors X and Y in a T-maze. Performance increases with shock intensity reaching a plateau at 60V. N=12 (15V), 12 (30V), 12 (60V) and 13 (90V). **c.** Experimental setup for *in vivo* calcium imaging. Flies are presented with electric shocks to the legs while neural activity is recorded in DANs expressing GCaMP6m. **d, f and h.** Mean ΔF/F0 calcium transients ± Standard Error of the Mean (SEM) measured from PPL1-γ1ped targeted by c061;MBGAL80 GAL4 (**d**), PPL1-γ2α’1 targeted by MB296B-GAL4 (**f**) and PPL1-α2α’2 targeted by MB058B-GAL4 (**h**) as flies were exposed to 0V, 15V, 30V, 45V, 60V and 90V shocks. **e, g and i.** Mean area under the curve ± Standard Deviation (SD) during the 1 min of 12 shocks for PPL1-γ1ped (**e**) (N=13, 15, 12, 13, 17 and 15), PPL1-γ2α’1 (N=13 for all groups) (**g**) and PPL1-α2α’2 (N=14, 12, 13, 10, 14 and 13) (**i**). **e.** PPL1-γ1ped DAN responses show a strong and significant correlation with shock intensity (Spearman correlation). **g.** PPL1-γ2α’1 DAN responses showing a strong and significant correlation with shock intensity (Spearman correlation). **i.** PPL1-α2α’2 DAN responses show no correlation with the shock intensity (Spearman correlation). Significant differences are calculated with a 1-way ANOVA and Dunnett’s multiple comparisons test or Kruskal-Wallis test and Dunn’s multiple comparisons test against 0V. Linear regression slopes (magenta) are tested against 0. Individual data points are displayed as dots. *P<0.05. ns: non-significant.

### Learning-induced plasticity at the KC to MBON-γ1ped>αβ and MBON-γ2α’1 junctions scales with shock intensity

During olfactory associative learning, dopamine from punishment or reward DANs targeting specific MB compartments drives synaptic depression between sparse odor-activated KCs and the corresponding MBONs in the relevant compartments, to promote appropriate learned behavior ^44,46,50,56,58–61,63^. We therefore used calcium imaging of MBON-γ1ped>αβ, MBON-γ2α’1 and MBON-α2sc odor responses after training to test whether PPL1-γ1ped, PPL1-γ2α’1 and PPL1-α2α’2 DANs might induce intensity-dependent plasticity of these KC-MBON connections when flies were trained with different voltages. Flies were trained under a 2-photon microscope (**Fig. 2a**; ^44^) using a differential conditioning paradigm where only the first of two odors was paired with different intensities of shock, the Condition Stimulus + (CS+) (**Fig. 2b**). We then immediately performed *in vivo* calcium imaging to measure odor-evoked responses after learning in MBON-γ1ped>αβ, MBON-γ2α’1 and MBON-α2sc, targeted by MB112C-GAL4, MB077B-GAL4 and MB080C-GAL4, respectively (**Supplementary Fig. 2a-c**). In line with previously published work ^44,50^, we found a robust depression of the CS+ relative to the CS-(unpaired odor) responses in MBON-γ1ped>αβ when flies were trained with 30V or higher intensity shock (**Fig. 2c-d**). Moreover, there was a high and significant correlation (r = 0.94) between the CS- CS+ response difference and the intensity of shock reinforcement (**Fig. 2i**). These results were recapitulated with the reciprocal pairing of odors as the CS+ and CS- (**Supplementary Fig. 2d, e and h**). MBON-γ2α’1 exhibited a significant difference in odor-evoked responses between CS+ and CS- when training was performed with 45V and higher voltages (**Fig. 2e-f**), again a result largely recapitulated in experiments with the opposite odors employed as CS+ and CS- (**Supplementary Fig. 2f, g and i**). A smaller difference was evident: there was a greater CS- increase after training when methylcyclohexanol (MCH) was the CS+ (**Fig. 2e-f**) compared to when 3-octanol (OCT) was the CS+ (**Supplementary Fig. 2f-g**). Nevertheless, both odor sequences showed a high and significant correlation (r = 1 and r = 0.99) between the CS- CS+ response difference and the intensity of shock (**Fig. 2j** and **Supplementary Fig. 2i**). These results, combined with our behavioral and PPL1 DAN recordings (**Fig. 1b** and **1d-g**), suggest that a threshold activation of PPL1-γ1ped (30V) and PPL1-γ2α’1 (45V) during learning is necessary to induce a behaviorally relevant memory trace in the KC to MBON-γ1ped>αβ and MBON-γ2α’1 connections. Similar experiments did not reveal learning-induced plasticity in MBON-α2sc odor responses (**Fig. 2g, h and k**), in agreement with the relative absence of PPL1-α2α’2 DAN shock responses irrespective of shock intensity (**Fig. 1h-i**).

**Fig. 2.**
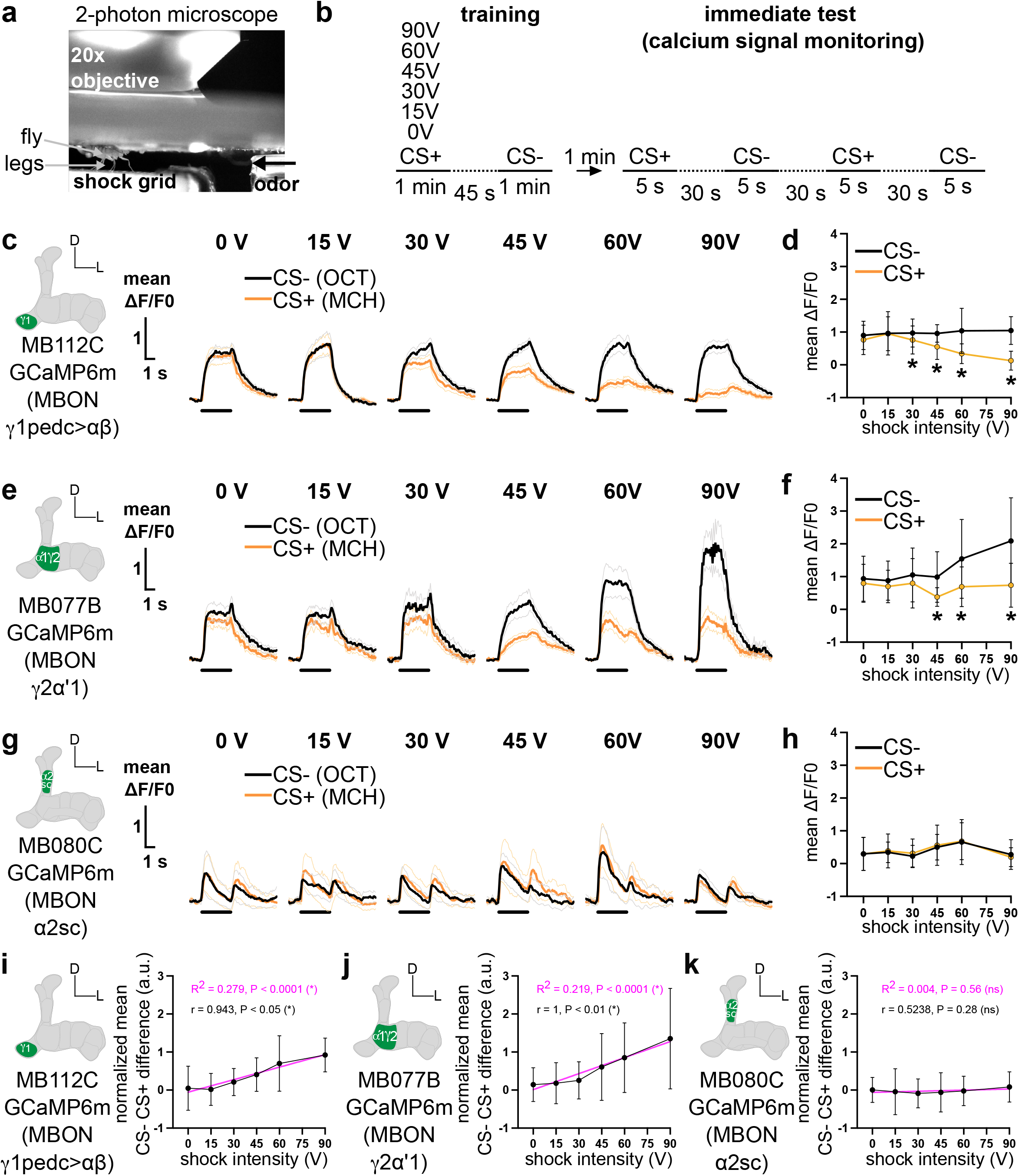
Aversive learning drives intensity-dependent plasticity of MBON-γ1ped >αβ, and MBON-γ2α’1 odor responses. **a.** Experimental setup. **b.** Protocol: Flies are trained to associate a 1 min odor presentation (CS+, here MCH) paired with electric shocks of 0V, 15V, 30V, 45V, 60V or 90V delivered to the legs, followed by a 45s presentation of fresh air and then another odor without shocks (CS-, here OCT). Immediately (1 min) after training neural activity is recorded in MBONs expressing GCaMP6m during 5 s presentations of the CS+ and CS-, performed twice and spaced by 30 s. **c, e and g**. Mean ΔF/F0 calcium transients ± SEM for the CS- (average of the two presentations, black) and for the CS+ (average of the two presentations, orange) immediately after training the CS+ with either 0V, 15V, 30V, 45V, 60V or 90V. MB112C is used to target MBON-γ1ped >αβ (**c**), MB077B for MBON-γ2α’1 (**e**) and MB080C for MBON-α2sc (**g**). **d, f and h.** Mean ΔF/F0 ± SD during the 5 s odor presentation for the CS- (black) and for the CS+ (orange) when paired with either 0V, 15V, 30V, 45V, 60V and 90V for MBON-γ1ped >αβ (N= 17, 14, 15, 16, 18 and 14) (**d**), MBON-γ2α’1 (N= 19, 16, 14, 15, 17 and 16) (**f**) and MBON-α2sc (N= 12, 11, 13, 11, 12 and 12) (**h**). Statistical tests are unpaired t-test (or Wilcoxon test) between the CS- and the CS+ for each voltage intensity. **i-k.** Normalized CS- CS+ difference ± SD and fitted linear regression for MBON-γ1ped >αβ (**i**), MBON-γ2α’1 (**j**) and MBON-α2sc (**k**) with a Pearson (r) (or Spearman) correlation. Linear regression slopes (magenta) are tested against 0. *P<0.05. ns: non-significant.

Our results show that both PPL1-γ1ped and PPL1-γ2α’1 DANs, and plasticity at MBON-γ1ped>αβ and MBON-γ2α’1 connections, correlate with the intensity of punishment during training. These data are consistent with the strength of the DAN response driving the changes in the CS+ and/or CS- odor responses in the corresponding MBONs. We therefore built a correlation matrix for all PPL1 DAN responses and all MBON CS- CS+ response differences using MCH as CS+ and OCT as CS- (**Supplementary Fig. 2j**) and the reciprocal odor combination (**Supplementary Fig. 2k**). We found a high and significant correlation between the DAN responses and MBON changes in the γ1ped and γ2α’1, but not α2α’2, compartments. These correlations also hold when OCT is the CS+ and MCH the CS- (**Supplementary Fig. 2k**). Overall, these data lead us to conclude that aversive memories of different electric shock intensity are written in a graded manner at the KC to MBON-γ1ped>αβ and MBON-γ2α’1 junctions, by the respective DANs whose activation scales with the intensity of electric shock. The combined intensity-dependent plasticity of odor-evoked responses in these MBONs will proportionally skew the overall MBON output to promote appropriate choices between the CS+ and CS- odors in the T-maze.

### Punishing and rewarding DANs are required to learn relative aversive value

Intensity dependent differential synaptic plasticity at multiple MB compartments could provide a substrate to compare and compute a relative aversive value between odors during learning, which can later be used during decision-making in the T-maze. To test this possibility and probe the underlying neural mechanisms, we used a relative aversive learning task. Flies were trained to associate three different odors (X, Y and Z) with different intensities of electric shock (0V, 60V and 30V, respectively) (**Supplementary Fig. 3a-b**; ^17,66^. The flies were then given either a relative choice between the Y_60_ vs Z_30_ odors, or an absolute choice between the X_0_ vs Y_60_ odors, in a T-maze. We first assessed the requirement of all PPL1 (**Fig. 3a**) or PAM (**Fig. 3b**) DANs during learning. The output of DANs was temporally blocked 30 min prior to and during training using expression of the dominant temperature-sensitive UAS-*Shibire*^ts1^ (*Shi*^ts1^) ^75^ at the restrictive temperature of 33°C. Flies were returned to permissive 23°C immediately after training, and 30 min later they were given a T-maze odor choice (**Fig. 3c**). As expected, blocking all aversive PPL1-DANs (MB504B-GAL4) during training impaired performance in both relative and absolute choice tests, compared to that of controls carrying only the GAL4 or *Shi*^ts1^ transgene (**Fig. 3d-e**). However, blocking PAM DANs (R58E02-GAL4) during training left absolute choices intact but abolished performance in the relative choice test (**Fig. 3d-e**; ^17^). Importantly, control experiments performed at permissive 23°C did not reveal significant differences between the groups (**Supplementary Fig. 3c-e**). Therefore, both reward and punishment coding DANs are necessary during learning to support appropriate relative choice.

**Fig. 3.**
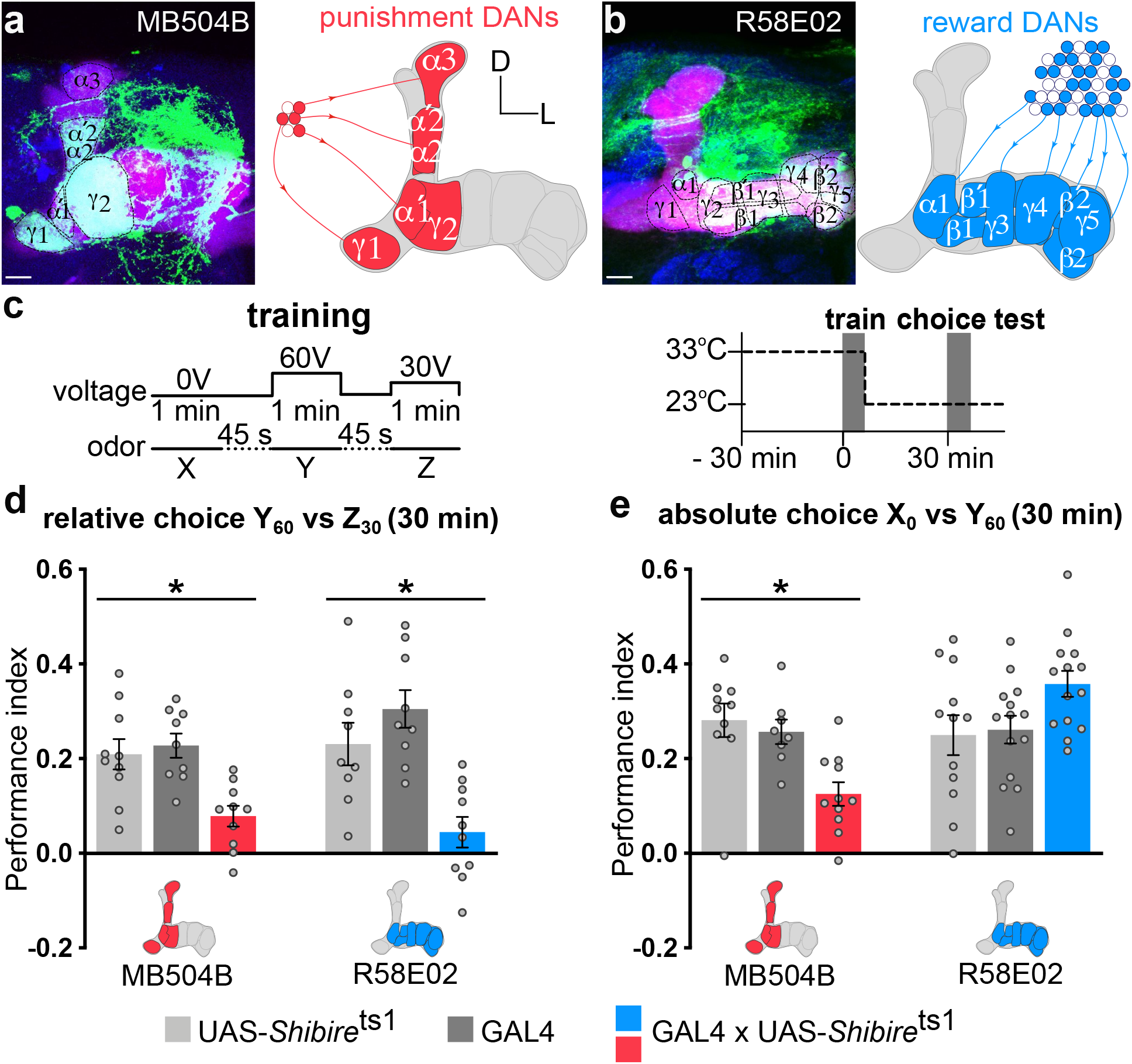
Aversive and reward DANs are required to learn relative aversive value whereas absolute value learning requires only aversive DANs. **a.** MB504-GAL4 driving UAS-mCD8::GFP labelling all PPL1 punishment DANs (schematized in red with the MB in gray). **b.** R58E02-GAL4 driving UAS-mCD8::GFP labeling most reward PAM DANs (schematized in blue with the MB in gray). In both a and b the right MB is co-labelled with 247-LexA::VP16-driven lexAop-rCD2::mRFP (magenta) and the whole brain with the presynaptic marker anti-Bruchpilot, nc82 (blue). Scale bar 10μm. **c.** Training protocol and temperature shifting procedure for *Shi*^ts1^ manipulations. **d.** Blocking output from MB504B- or R58E02-GAL4 DANs during training impaired 30 min relative choice compared to that of relevant controls (both 1-way ANOVA, N=9-10, P<0.05). **e.** Blocking output from MB504B- but not R58E02-GAL4 during training impaired 30 min absolute choice compared to that of relevant controls (Kruskal-Wallis, N=8-11, P<0.05 and 1-way ANOVA, N=12-14, P>0.05, respectively). Data are mean ± SEM. Individual data points are displayed as dots. *<P<0.05.

### PPL1-γ1ped, PPL1-γ2α’1 and PPL1-α2α’2 aversive DANs are required to learn relative aversive value

Prior work has defined the DANs conveying aversive reinforcement as the 4 different subtypes from the PPL1 cluster and a few neurons from the PAM cluster (**Fig. 4a**; ^36^). We tested which of the PPL1-γ1ped (c061;MBGAL80; **Supplementary Fig. 4a**), PPL1-γ2α’1 (MB296B-GAL4; **Supplementary Fig. 4b**), PPL1-α2α’2 (MB058B-GAL4; **Supplementary Fig. 4c**), PPL1-α3 (MB630B-GAL4; **Supplementary Fig. 4d**) and PAM-β2β’2a (using NP5272-GAL4; **Supplementary Fig. 4e** and MB301B-GAL4; **Supplementary Fig. 4f**) DAN subtypes were critical to learn relative and absolute aversive value. We did not test PAM-γ3^45^. We again used *Shi*^ts1^ to block their output during learning and assessed their requirement in a relative or absolute odor choice test 30 min after training (**Fig. 4b**). Blocking PPL1-γ1ped DANs during learning reduced both relative and absolute choices (**Fig. 4c-d**). In contrast, blocking PPL1-γ2α’1 or PPL1-α2α’2, but not PPL1-α3 or PAM-β2β’2a, during learning left absolute choices unchanged but abolished relative choices (**Fig. 4c-d** and **Supplementary Fig. 4g-i**). No defects in performance were apparent when the entire experiment was performed at permissive 23°C (**Supplementary Fig. 4j-l**). Together, these results reveal PPL1-γ1ped as critical for signaling absolute and relative aversive value while PPL1-γ2α’1 and PPL1-α2α’2 are only required to learn relative aversive value.

**Fig. 4.**
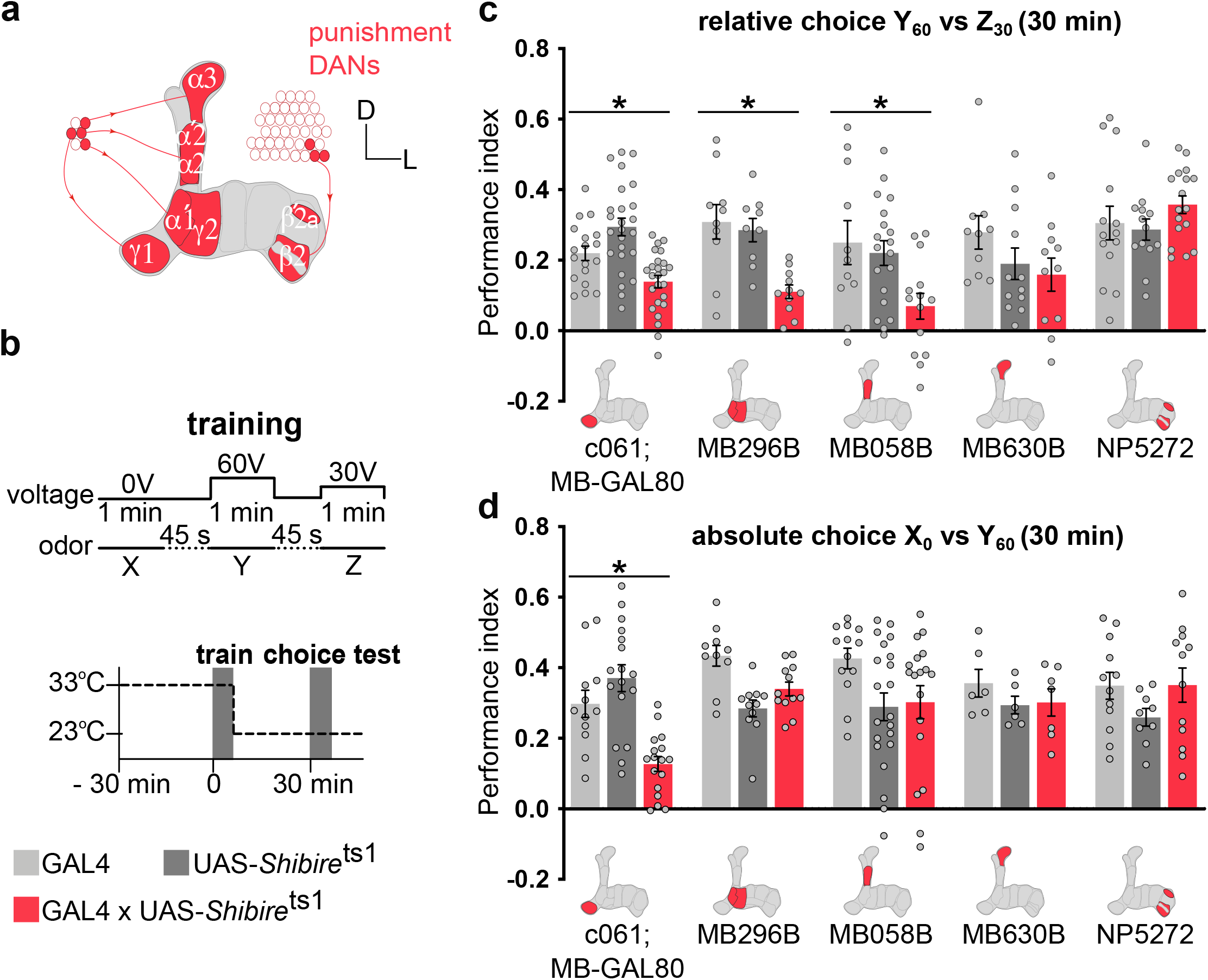
PPL1-γ1ped, PPL1-γ2α’1 and PPL1-α2α’2 DANs are required to learn relative aversive value while only PPL1-γ1ped is necessary to learn absolute aversive value. **a.** Schematic of MB innervation by aversively reinforcing DANs. **b.** Training protocol and temperature shifting procedure for *Shi*^ts1^ manipulations. **c.** Blocking output from c061;MBGAL80- (PPL1-γ1ped), MB296B- (PPL1-γ2α’1) and MB058B- (PPL1-α2α’2) but not MB630B- (PPL1-α3) or NP5272- (PAM-β2β’2a) GAL4 neurons during training impaired 30 min relative choice compared to that of relevant controls (1-way ANOVA, N=18-25, P<0.05; 1-way ANOVA, N=9-11, P<0.05; 1-way ANOVA, N=11-19, P<0.05; Kruskal-Wallis, N=10-12, P>0.05; 1-way ANOVA, N=13-17, P>0.05 respectively). **d.** Blocking output from c061;MBGAL80- (PPL1-γ1ped) but not from MB296B- (PPL1-γ2α’1), MB058B- (PPL1- α2α’2), MB630B- (PPL1-α3) or NP5272- (PAM-β2β’2a) GAL4 neurons during training impaired 30 min absolute choice compared to that of relevant controls (1-way ANOVA, N=12-17, P<0.05; Kruskal-Wallis, N=10-12, P>0.05; 1-way ANOVA, N=13-21, P>0.05; Kruskal-Wallis, N=6-7, P>0.05; 1-way ANOVA, N=9-12, P>0.05 respectively). Data are mean ± SEM. Individual data points are displayed as dots. *<P<0.05.

### PAM-β’2aγ5n rewarding DANs are required to learn relative value

We next tested which specific rewarding PAM DANs (**Fig. 5a**) were required to learn relative aversive value. We employed the same protocol as before in **Fig. 3** and **Fig. 4** (**Fig. 5b**) and used *Shi*^ts1^ to block the output of discrete subpopulations of rewarding PAM DANs during learning. Flies were then given a relative odor choice 30 min after training (**Fig. 5b**). We used 0104-GAL4, which labels DANs innervating the β2sc, β’2a, β’2mp, γ4 and γ5b (broad) MB compartments (^17,40,41,76^; **Supplementary Fig. 5a**), R56H09-GAL4 which labels DANs innervating β’2a, β’2mp, γ4 and γ5n (narrow) (^41^; **Supplementary Fig. 5b**) and 0279-GAL4 labeling DANs in β1 and β2 (^17,41^; **Supplementary Fig. 5c**). Blocking 0104-GAL4 and R56H09-GAL4, but not 0279-GAL4, DANs only during learning significantly impaired relative choice test performance (**Fig. 5c**). Importantly, control experiments performed at permissive 23°C did not reveal significant differences between the relevant groups (**Supplementary Fig. 6a-b**). In addition, these subsets of reward DANs were not required during learning when flies were given an absolute choice, suggesting that both olfactory and shock perception were unaffected by the manipulations (**Supplementary Fig. 6c-d**). We next assessed whether subsets of reward DANs included in R58E02-GAL4 (most reward DANs), but not in R56H09-GAL4, are involved in relative aversive value coding. We generated a R56H09-GAL80 line and combined it with R58E02-GAL4 to restrict the *Shi*^ts1^ block to the remaining α1, β1, β2sc, γ3 and β’1 DANs (**Supplementary Fig. 5d**). These remaining DANs were not required during learning for performance in either relative (**Fig. 5c**) or absolute (**Supplementary Fig. 6c-d**) choice tests.

**Fig. 5.**
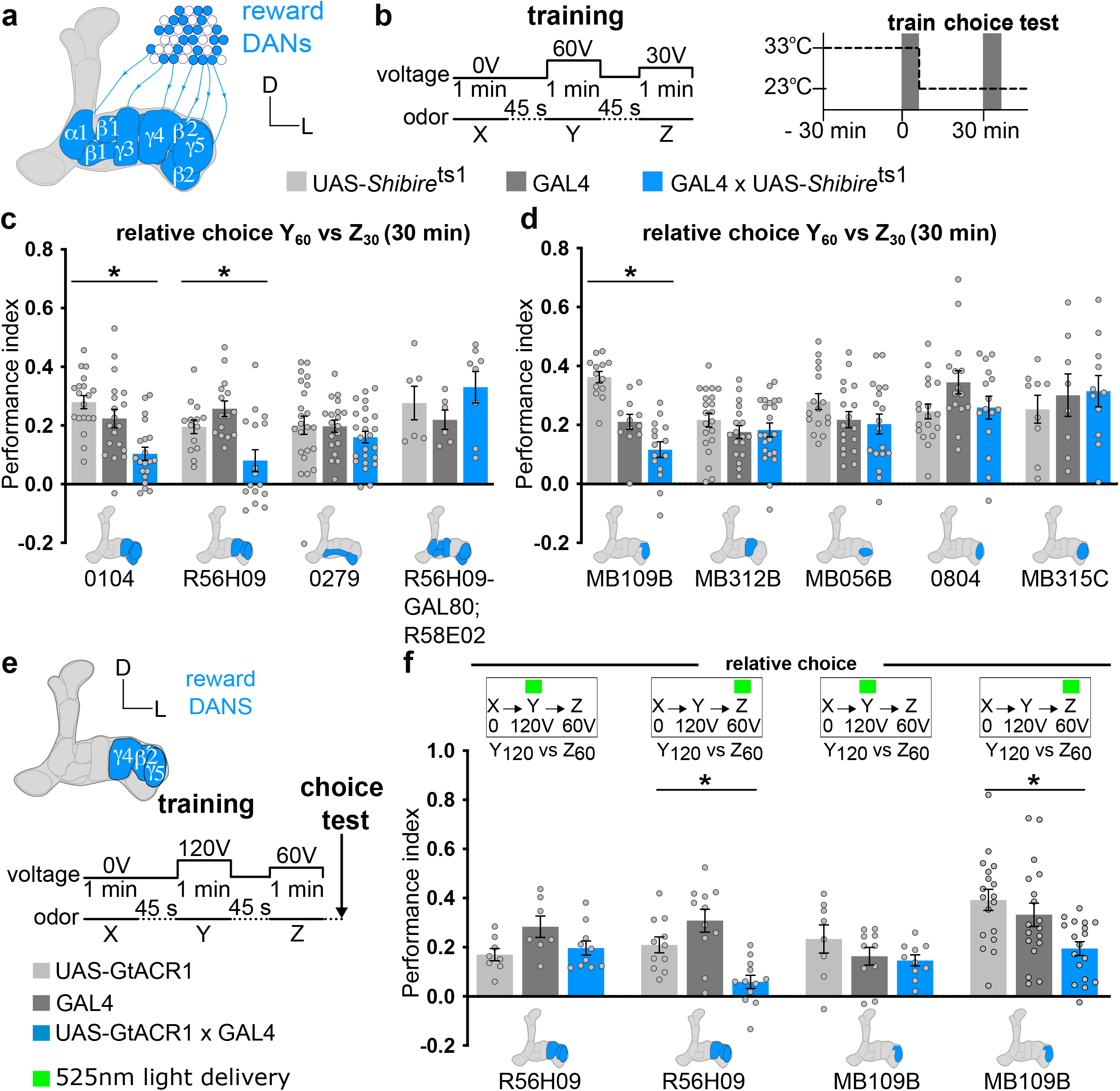
PAM-β’2aγ5n DANs are required to learn relative aversive value. **a.** Schematic of MB innervation of reward DANs. **b.** Training protocol and temperature shifting procedure for *Shi*^ts1^ manipulations. **c.** Blocking output from 0104- and R56H09- but not from 0279- or R56H09-GAL80;R58E02-GAL4 targeted PAM DANs during training impaired 30 min relative choice compared to that of relevant controls (1-way ANOVA, N=19- 20, P<0.05; 1-way ANOVA, N=14-16, P<0.05; 1-way ANOVA, N=19-24, P>0.05; Kruskal-Wallis, N=6-8, P>0.05; respectively). **d.** Blocking output from MB109B- but not of MB312B-, MB056B-, 0804-or MB315C-GAL4 targeted PAM DANs during training impaired 30 min relative choice compared to that of relevant controls (1-way ANOVA, N=12-15, P<0.05; 1- way ANOVA, N=18-22, P>0.05; 1-way ANOVA, N=16-18, P>0.05; 1-way ANOVA, N=15-18, P>0.05; 1-way ANOVA, N=9-11, P>0.05; respectively). **e.** Schematic of reward DAN innervation for R56H09- and MB109-GAL4 and training protocol. **f.** Inhibiting reward DANs with a 525nm green light exposure only during the last odor Z + 60V association impaired immediate relative choice (1-way ANOVA, N=11-12, P<0.05 for R56H09-GAL4 and 1-way ANOVA, N=18, P<0.05 for MB109B-GAL4), but not when inhibited during the odor Y + 120V association (1-way ANOVA, N=6-10, P>0.05 for R56H09-GAL4 and 1-way ANOVA, N=8-10, P>0.05 for MB109B-GAL4). Data are mean ± SEM. Individual data points are displayed as dots. *<P<0.05.

We then assessed the role of the individual subtypes of rewarding DANs labeled by R56H09-GAL4 in learning relative aversive value (**Fig. 5d**). We used the following specific GAL4 driver lines: PAM-β’2aγ5n (MB109B-GAL4; **Supplementary Fig. 5e**), PAM-γ4 and γ4>γ1γ2 (MB312B-GAL4; **Supplementary Fig. 5f**), PAM-β’2mp (MB056B-GAL4; **Supplementary Fig. 5g**), PAM-β2sc and γ5n (0804-GAL4; **Supplementary Fig. 5h**) and PAM-γ5b (MB315C-GAL4; **Supplementary Fig. 5i**). Blocking output with *Shi*^ts1^ only revealed a specific requirement for the β’2aγ5n reward DANs to learn relative aversive value (**Fig. 5d**). Control experiments at permissive 23°C showed no statistical differences between the groups (**Supplementary Fig. 6a-b**). In addition, these reward DANs were not required during learning for absolute choice (**Supplementary Fig. 6e**). We also tested a second GAL4 line (MB087C-GAL4) which labels fewer DANs innervating the β’2a and γ5n compartments, than are labeled in MB109B-GAL4 (**Supplementary Fig. 5e, j and k**). Blocking the output of MB087C-GAL4 neurons during learning did not alter relative or absolute choices (**Supplementary Fig. 6f-g**). Since all flies exhibited normal absolute choices, we conclude that both olfactory and shock perception remained intact from the thermogenetic manipulations during training. Together, these results show that particular (or a critical number of) PAM-β’2aγ5n reward DANs are necessary to learn relative aversive value and therefore enable appropriate relative-value based choice.

To signal relative aversive value (‘less bad’ or ‘better than’) during learning, the PAM-β’2aγ5n should only be necessary during the last lesser 30V experience, when a comparison can be made between current 30V and previous 60V aversive experience. However, the temporal resolution afforded by the thermogenetic *Shi*^ts1^ is not adequate to easily test this hypothesis. We therefore used optogenetic inhibition to control rewarding DAN activity during specific segments of the learning session. In addition, classical shock training chambers (as used in all prior experiments) contain copper wires, which decreases light penetrance and impedes the use of optogenetics. We therefore designed a transparent shock grid made of indium tin-oxide coated on plastic that could be wrapped inside the training chamber (inspired by a prior apparatus for electric shock-reinforced visual learning ^77^). Due to the higher electrical resistance of this material, we altered the training voltages to achieve a similar efficiency of training. Thus, the three different odors (X, Y and Z) were respectively paired with 0V, 120V and 60V shocks (**Fig. 5e**). During learning, we delivered light with a spectrum peaking at 525nm to the flies to inhibit the targeted reward DANs (R56H09-GAL4 or MB109B-GAL4) expressing the green light-sensitive anion channelrhodopsin GtACR1 ^78^. Reward DANs were photo-silenced only during the second or the third odor/shock association and 1 min later flies were tested in a relative (Y_120_ vs Z_60_) or absolute (X_0_ vs Y_120_ or X_0_ vs Z_60_) odor choice. Strikingly, reward DANs were only required during the last training segment (Z + 60V), being dispensable during the second training segment (Y + 120V), to enable an appropriate relative choice (**Fig. 5f**). Control experiments without light stimulation showed no differences in the 1 min relative choice test (**Supplementary Fig. 7a-b**). Moreover, reward DANs were not needed during the second or third odor/shock association to enable appropriate absolute (X_0_ vs Y_120_ or X_0_ vs Z_60_) choices (**Supplementary Fig. 7c**). These results suggest that the PAM-β’2aγ5n reward DANs are crucial to learn a relative ‘better-than’ aversive value between current and previous aversive experiences.

### PAM-β’2aγ5n DANs integrate MBON-γ2α’1 input to signal relative aversive value

To potentially visualize a relative ‘better-than’ reward value signal, we expressed GCaMP6m and recorded calcium transients from PAM-β’2aγ5n DANs while training flies with the relative aversive value protocol under a 2-photon microscope (**Fig. 6a-c**). We used a mock training protocol as a control, where the same odor presentations were given as in the relative aversive training condition, but no shocks were delivered. We analyzed the data by dividing the recordings made during each 1 min X, Y and Z odor presentation into four consecutive 15 s segments. No differences were apparent in PAM-β’2aγ5n DAN calcium responses in trained flies (blue) compared to those of the mock controls (black) during the first odor X, or second odor Y with high 60V shocks, presentations (**Fig. 6d**). However, we observed a significant elevation of PAM-β’2aγ5n responses within the first quarter of the recordings when the last odor Z was presented with lesser 30V shocks (**Fig. 6d**). In addition, we noted that the elevated PAM-β’2aγ5n DAN responses occurred after the first of the 30V shocks was delivered. Combined with our behavioral results in **Fig. 5f**, these data are consistent with a model in which PAM-β’2aγ5n DANs compare previous and current aversive experience to signal a relative aversive ‘better than’ reward signal during learning, which subsequently allows the flies to make an appropriate relative value-based odor choice.

**Fig. 6.**
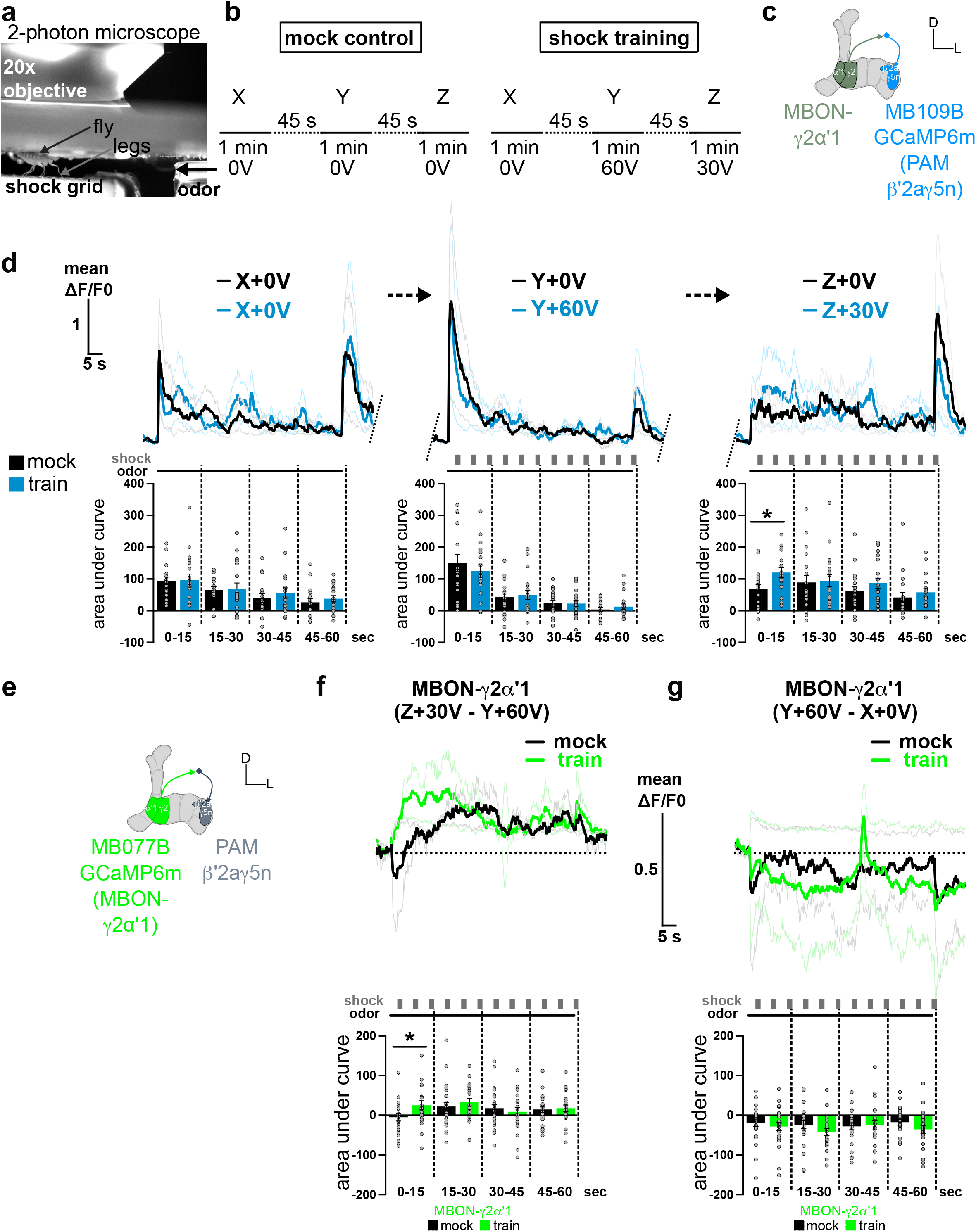
PAM-β’2aγ5n DANs integrate MBON-γ2α’1 input to signal ‘better than’ value during relative aversive learning. **a.** Experimental setup. **b.** Control mock protocol: the odors X, Y and Z are presented alone. For shock training: the odor X is unpaired, odor Y is paired with 60V, and odor Z with 30V. **c.** Schematic of the MB innervation of MBON-γ2α’1 and recorded PAM-β’2aγ5n (MB109B-GAL4 driving UAS-GCaMP6m) and their synaptic contact. **d.** PAM-β’2aγ5n DAN calcium traces and mean area under curve quantifications for each 15 sec quarter of every 1 min odor presentation for the mock (black, N=19) and shock trained (blue, N=19) protocols. A significant increase in the trained protocol odor-responses were only observed during the first quarter of the odor Z + 30V presentation compared to that of the mock control recordings (unpaired t test; P<0.05). All other comparisons with unpaired t-test or Mann-Whitney are non-significant (P>0.05). **e.** Schematic of MB innervation of recorded MBON- γ2α’1 (MB077B-GAL4 driving UAS-GCaMP6m) and PAM-β’2aγ5n and there synaptic contact. **f.** Mean subtracted Z+30V – Y+60V calcium traces and quantifications of mean area under the curve of MBON-γ2α’1 during the trained (N=23) and mock (N=26) protocols. A significant increase was only found during the first quarter of the odor presentation period in the Z+30V – Y+60V subtracted responses of the trained group compared to that of the mock group (unpaired t test; P<0.05). All other comparisons are non-significant (unpaired t or Mann-Whitney tests; P>0.05). **g.** Mean subtracted Y+60V – X+0V calcium traces and mean area under curve of MBON-γ2α’1 during the train (N=23) and mock (N=26) protocols. We found no significant differences during the odor presentation subtraction Y+60V – X+0V (unpaired t and Mann-Whitney tests; P>0.05). Data are mean ± SEM. Individual data points are displayed as dots. * P<0.05.

To understand how the PAM-β’2aγ5n DANs integrate and compare previous and current aversive experience to signal relative aversive value we considered their synaptic input connectivity in the MB network. Strikingly, PAM-β’2aγ5n DANs receive direct synaptic input from MBON-γ1ped>αβ ^53^ and MBON-γ2α’1 ^46,53^, both of whose odor-evoked responses are modified by aversive learning (**Fig. 2** and **Supplementary Fig. 2**). MBON-γ5β’2a also connects to PAM-β’2aγ5n DANs in recurrent connections ^76^ and their odor-evoked responses are potentiated by aversive learning ^44,50,59,60^. We therefore made independent recordings from these three MBONs during the relative aversive training protocol (**Supplementary Fig. 8**). No obvious differences in the odor-evoked responses of these MBONs were apparent during each of the three odor sequences (X + 0V, Y + 60V and Z + 30V), as compared to flies presented with the mock training protocol (**Supplementary Fig. 8**). Since our model proposes that PAM-β’2aγ5n DANs might compare current and previous aversive experiences coded in each MBON during the last odor Z + 30V association, we subtracted the (previous) odor-evoked responses during the odor Y + 60V association from those during the (current) odor Z + 30V association. This data analysis uncovered a significant increase in the differential responses of MBON-γ2α’1 during the first quarter (the first 3 shocks) of the odor/shock training sessions, as compared to those in flies subjected to the mock training protocol (**Fig. 6e-f**). Again, and as with PAM-β’2aγ5n DANs, this differential increased response seems evident after the first of the 30V shocks was delivered. In contrast, subtractive analyses did not reveal any differences between present (Z + 30V) and prior (Y + 60V) responses for MBON-γ1ped>αβ or MBON-γ5β’2a (**Supplementary Fig. 9a-d**). In addition, no statistical differences were evident for any MBON when we subtracted the odor-evoked response during the odor Y + 60V association from those during the first unpaired odor X + 0V presentation (**Fig. 6g** and **Supplementary 9e-h**). We therefore propose that comparing current and previous aversive experiences requires additional cholinergic input from MBON-γ2α’1 during the odor Z + 30V to trigger the PAM-β’2aγ5n DANs, which signal a ‘better than’ relative aversive value to the lesser aversive olfactory experience.

## Discussion

Our study addresses how animals assign absolute aversive value during learning and how they compare and ascribe relative aversive value information to consecutive negative experiences, so that they can making appropriate value-based decisions afterwards. Using the fruit fly *Drosophila* permitted a cellular resolution view of how the interaction between the appetitive and aversive DAN systems, within the neural network of the MB, is at the heart of the mechanistic underpinnings that compute a relative aversive value during associative olfactory learning.

### Aversive experience is uniquely coded at the KC-MBON-γ1ped>αβ and -MBON-γ2α’1 junctions

PPL1 DANs reinforce a range of aversive memories with differing strength and persistence ^34, 36–38,55,58,79,80^. Our data provide new insight into the functional diversity of these anatomically discrete DANs (**Fig. 1, 4, 5**, and **Supplementary Fig. 1, 3 and 4**). We found that the individual aversively reinforcing PPL1-γ1ped, PPL1-γ2α’1 and PPL1-α2α’2 DANs exhibit different intensity response profiles when flies were exposed to a series of shock voltages ^71,72^. Importantly, the strength of their responses to electric shocks strongly correlated with the magnitude of plasticity of the odor-evoked responsiveness of their corresponding MBONs after differential conditioning (the first of two odors being paired with electric shocks of different intensities). These results indicate that absolute aversive value is assigned to odors in different ways in the γ1ped, γ2α’1 and α2α’2 MB compartments, consistent with the conclusion of a prior study that artificially-activated the individual DANs^58^. Of note, we did not observe significant shock responses in PPL1-α2α’2 DANs (**Fig. 1h-i**). These results are also in accordance with the absence of odor-evoked changes in the corresponding MBON-α2sc after training (**Fig. 2g-h**) and a lack of reinforcing properties found by pairing artificial activation of these neurons alone with an odor ^37^ (but see ^58^). We observed that the magnitude of the MBON-γ1ped>αβ CS+ responses depression is tightly correlated to the voltage applied to the flies, and the resulting activity of the PPL1-γ1ped DANs. In other words, the stronger the aversive experience, the greater the PPL1-γ1ped DAN driven depression of the CS+-evoked response of MBON-γ1ped>αβ. In comparison, the responses of the CS- odor appeared unaffected by absolute aversive conditioning (**Fig. 2c-d**). Feedforward GABAergic inhibition from MBON-γ1ped>αβ to the primary axon of MBON-γ5β’2a ^44,50^ is therefore reduced in a graded manner by aversive conditioning. MBON-γ5β’2a should therefore display a proportional increase in its CS+-evoked response to drive learned avoidance behavior ^59,60^. Our experiments uncovered a very different effect of absolute aversive conditioning at the MBON-γ2α’1 junction. Although the PPL1-γ2α’1 DANs were significantly triggered by shocks ≥30V, their responses were comparable at all voltages between 30 and 90V (**Fig. 1f-g**). Moreover, aversive conditioning did not significantly depress the CS+ responses of MBON-γ2α’1 (**Fig. 2e-f** and **Supplementary Fig. 2f-g**). Instead, we observed that the responses of the CS- odor were increased and the CS- CS+ differential response were correlated with the intensity of the shocks applied (**Supplementary Fig. 2**) (these results differ from those of ^61^, perhaps due to a different response comparison). Our data therefore suggest that any odor that follows the CS+ with ≥45V presentation during training gains the capacity to drive more activity in the cholinergic MBON-γ2α’1. In addition, our recordings indicate that the more aversive the first experience is the stronger the cholinergic MBON-γ2α’1 activity will be to the subsequent experience. These data support the key involvement of MBON-γ2α’1 in the coding of relative aversive value.

### MBON-γ2α’1 input to PAM β’2aγ5n DANs provides a ‘better than’ reward signal during relative aversive training

We found that output from the PAM β’2aγ5n DANs was critical during the odor Z + 30V presentation for relative aversive learning (**Fig. 5**). These DANs receive direct excitatory cholinergic input from MBON-γ2α’1 ^46,53^ and we propose that the strength of this excitation is key for the flies to assign a ‘better than’ reward value to the lesser of the two aversive experiences. As mentioned above, when odor Y is paired with 60V shock in a differential conditioning assay the CS- responses of MBON-γ2α’1 become elevated. This means that when Y + 60V is followed by Z + 30V, the Z odor will more strongly drive MBON-γ2α’1 and as a result will activate the PAM β’2aγ5n DANs (**Fig. 6d**). In effect, any odor that follows a Y + 60V experience is predisposed to be judged as ‘better than’, unless it is itself accompanied by 60V or greater voltage. Our analyses that subtracted MBON-γ2α’1 odor-evoked responses are entirely consistent with this model. Odor driven activity of MBON-γ2α’1 is greater during the first period of the following Z + 30V experience than during the same period (just after the first shock) of the prior Y + 60V experience (**Fig. 6f**). Critically, this is also the time period during which we observe an elevation of PAM β’2aγ5n DAN activity (**Fig. 6d**). We speculate that the first of the 30V shocks somehow further releases the PAM β’2aγ5n DAN activity to be fully driven by MBON-γ2α’1, perhaps as a release of feedforward inhibition in the MBON-γ1ped>αβ to MBON-γ5β’2a to PAM β’2aγ5n DAN pathway. Our results and proposed models of PAM-β’2aγ5n DANs providing a ‘better than’ reward signal are in accordance with previous reports that PAM-β’2aγ5n activation provides appetitive reinforcement ^37,41,46,81^.

### Are there limits to comparable aversive memories?

Individual PPL1-DAN subtypes have different thresholds for activation and intensity-dependent plasticity in their corresponding MBON junctions have similar thresholds. We noted that these thresholds seem reflected in the range of comparisons that flies can make in a relative choice between different aversive memories ^66^, which point towards a threshold and a difference between voltages of 30V as being optimal to efficiently estimate a relative difference. In our recordings 30V was the threshold for observing shock-evoked responses in PPL1-γ1ped and PPL1-γ2α’1, but did not trigger PPL1-α2α’2. In addition, 30V produced significant plasticity of MBON-γ1ped>αβ odor responses but plasticity was not evident in MBON-γ2α’1 responses until 45V. Thus, perhaps every odor paired with voltage of ≤30V is considered to be ‘not so bad’ because it only depresses the GABAergic MBON-γ1ped>αβ responses and not the cholinergic MBON-γ2α’1 responses, thereby leaving CS+ odor driven excitation of PAM β’2aγ5n DANs from these MBONs. Although flies can differentiate between stronger aversive memories such as 90V vs 60V their relative choice performances are less good than 60V vs 30V ^66^. While we did not observe significant shock responses for PPL1-α2α’2 DANs, we found a role for these neurons during learning of relative aversive value (**Fig. 4**). MBON-γ1ped>αβ is GABAergic and is connected to PPL1-α2α’2 DANs ^38,53^. It is therefore possible that repeated pairing of odor Y + 60V electric shocks (or anything above their threshold) during relative training induces enough CS+ -evoked depression at MBON-γ1ped>αβ to release inhibition in PPL1-α2α’2 DANs while pairing the odor Z with 30V shocks ^82^. The resulting plasticity in MBON-α2sc could explain the requirement of αβ surface and αβ core KCs during a relative choice Y_60_ vs Z_30_ ^17^.

### Which PAM-β’2aγ5n DANs are responsible for ‘better than’ valuation?

Our results (**Fig. 5**) narrow down ‘better than’ valuation to a population of reward DANs in the PAM cluster that innervate the β’2a and γ5 MB compartments. The crucial MB109B-GAL4 line has been previously described as being restricted to PAM-β’2a DANs ^37,46,81,83^. However, this split-GAL4 line also shows clear expression in neurons that innervate a narrow part of the γ5 compartment (**Supplementary Fig. 5e**; ^84^). For this reason, we have named these neurons as PAM-β’2aγ5n throughout. Recent ultrastructural analyses have established that there is additional structural heterogeneity of both PAM-β’2a and PAM-γ5 DANs ^53,76^. Importantly, only some of these DANs receive input from MBON-γ2α’1 ^53,76^. Considering the cell body count, the commissure crossing ^76^ between the MB horizontal lobes, and a requirement for MBON-γ2α’1 input of neurons in MB109B-GAL4, it appears that this line includes three DAN types: PAM-γ5 dorsal dendrite (PAM01fb) (those that receive feedback from MBON-γ5β’2a, MBON01 ^76^), PAM-β’2a middle and lower commissure (PAM02), and also the recently identified PAM-γ5β’2a (PAM15) that has processes in both γ5 and β’2a ^53^. MB087C-GAL4 expresses in DANs that innervate β’2a and γ5 but it labels fewer neurons than MB109B-GAL4 (**Supplementary Fig. 5j-k**) and blocking the output of these neurons had no consequence for learning relative aversive value (**Supplementary Fig. 6f-g**). It is possible that MB109B-GAL4 does not express in enough of the relevant DANs, or it is missing expression in a crucial subtype receiving MBON-γ2α’1 inputs. Further investigation, and probably refinement of available tools, will be needed to disentangle the exact PAM-β’2aγ5n subtypes that are critical for coding relative aversive value.

### Relative positive evaluation of aversive experiences

Learning relative aversive value requires an interplay between aversively reinforcing PPL1 DANs modulating KC-MBON connections which provide feedforward and recurrent feedback input that determines the activity of specific subtypes of rewarding PAM DANs. These results support long-held ^85,86^ and recent ^44–46,54,87,88^ models in both vertebrates and invertebrates suggesting that learning requires critical interactions between appetitive and aversive reinforcement systems. In the fly, and likely also mammals, this process relies on opposing populations of DANs providing predictive signals needed to compare current with previous experience to assign (and update) both absolute and relative value to stimuli during learning. For instance, aversive memory extinction and reversal learning require the reward system in both vertebrates and invertebrates ^44,88–90^. In all these cases, stimuli that represent the absence of a punishment are rewarded ^44,46,91^. In humans, the ventral striatum, targeted by numerous DA inputs from the ventral tegmental area (VTA) providing rewarding information, is essential to compare aversive experiences of different intensities ^16,27^. In the orbitofrontal cortex, relative coding of aversive (but also appetitive) experiences seem to require overlapping neuronal ensembles to select a preferred option and promote appropriate economical decisions in a specific spatial and temporal context ^12,18,92,93^. In the dopaminergic system reward is also computed in a relative manner to broadcast value signals in different brain regions ^6^. These DANs from the VTA and substantia nigra compute a prediction error to signal positive, but also negative, value (for reviews see ^23,24^). A similar value prediction error calculation has not yet been demonstrated experimentally in the fly ^72,94^. Instead, results from several studies in the fly suggest that errors are registered in the MB network by the action of DANs that signal the opposing value ^44,46,95,96^. Our experiments here suggest that a similar interplay between opposing populations of DANs, and plasticity at different MBON junctions in the MB network, permits computation of relative aversive value (or difference) between a previous and a new aversive experience. Combined with previous work ^44,45,54,55,97^ and current computational models ^70,98^, our data provide key features of how a heterogeneous DAN system can “pre-compute” a relative value during learning ^5^ that facilitates future value-based decisions.

## METHODS

### Resource availability

Further information and requests for resources and reagents should be directed to, and will be fulfilled by, the lead contact, Emmanuel Perisse.

### Materials availability

All original reagents presented in this study are available from the Lead Contact upon request.

### Data and code availability

All code used in this paper with Fiji or MATLAB will be shared by the Lead Contact upon request.

## EXPERIMENTAL MODEL AND SUBJECT DETAILS

### Animals

Fly stocks were cultured at 21°C on standard cornmeal food which was made with 40L of tap water, 280g of agar, 1.5kg of yeast, 2kg of corn flour, 2.8kg of Dextrose, 120g of Tegosept and 460mL of Ethanol. The R56H09-GAL80 construct was made by amplifying the R56H09 enhancer region from the genomic DNA of R56H09-Gal4 flies (Pfeiffer et al., 2010) using forward primer 5’-CACCGGCTACCACACCCAGCGTGCAACAG-3’ and reverse primer 5’- CCTCCTTGATCAGGCGCGAACAGGT-3’. The PCR product was cloned into pENTR/D-TOPO vector (pENTR™ Directional TOPO® Cloning Kits, Invitrogen). Then the enhancer fragment was put into pBPGal80Uw-6 plasmid (Addgene) using Gateway cloning system (Gateway™ LR Clonase™ II Enzyme Mix, Invitrogen). The R56H09-GAL80 plasmid was inserted into the attp40 landing site by site-specific integration (Bestgene). Genotypes and sources of the fly lines used in this study are denoted in **Table S1**.

## METHOD DETAILS

### Behavioral Analyses

Behavioral experiments were performed as previously described ^17^ with minor modifications. For all experiments, groups of ∼100 5-10 day old flies were housed for 18–24 h before training in a 25 ml vial containing standard cornmeal food and a 20 x 60 mm piece of filter paper at 23°C and 65% relative humidity and a 12 h/12 h light-dark cycle (except for optogenetic experiments for which flies were raised in the dark prior to the experiment). Electric shocks were delivered with a Grass S48 Square Pulse Stimulator (Grass Technology) for behavioral experiments or a combination of DS2A (voltage stimulator) and DG2A (trained/delay generator) (Digitimer Ltd, UK) for *in vivo* calcium imaging experiments. For experiments in Fig. 1a-b and all experiments using the thermogenetic tool *Shi*^ts1^, we used a training tube with copper wires covering the inside of the tube. For the experiments in Fig. 1a-b, flies were trained at 23°C and 65% relative humidity as follows: 1 min odor X with 12 x 1.5 s shocks (at 0.2 Hz) of either 15V, 30V, 60V or 90V, followed by 45 s air, and 1 min odor Y without reinforcement.

For the *Shi*^ts1^ experiments at restrictive temperature, flies were transferred 30 min prior to and during training into a behavioral room at 33°C and 65% relative humidity. Flies were trained as follows: 1 min odor X unpaired, followed by 45 s air, 1 min of odor Y with 12 x 1.5 s 60V shocks (at 0.2 Hz), followed by 45 s air, and 1 min odor Z with 12 x 1.5 s 30V shocks (at 0.2 Hz).

For the optogenetic experiments, we designed a transparent shock grid made of a 75 mm x 50 mm polyethylene terephthalate (PET) coated with a conductive indium tin oxide (ITO) 175μm film of 80 Ohms / Sq (Diamond Coatings Ltd., UK). A grid was laser-etched onto the ITO film in order to insulate the positive and negative electrodes (lanes in the grid were 0.5 mm spaced by 0.3 mm apart). Flies were trained at 23°C and 65% relative humidity as follows: 1 min odor X unpaired, followed by 45 s air, 1 min of odor Y with 12 x 1.5 s 120V shocks (at 0.2 Hz), followed by 45 s air, and 1 min odor Z with 12 x 1.5 s 60V shocks (at 0.2 Hz). The GtACR1 optogenetic stimulation was provided by 4 green LEDs at 101.85μW/mm^2^ (λ = 525nm; Prolight Opto, PM2B-3LxE-SD, Technology Corporation, Taiwan) constantly illuminating the entire training tube during the presentation of odor Y or Z.

Memory performance (immediately or 30 min after conditioning) was tested in the dark by allowing the flies to choose for 2 min between two of the odors presented during training: X vs Y for Fig. 1a-b, relative choice Y vs Z and absolute choices X vs Y or X vs Z for all other behavioral experiments. A Performance Index (PI) was calculated as the number of flies making the correct choice minus the number of flies making the wrong choice, divided by the total number of flies in each experiment. A single PI value is the average score from flies of the identical genotype tested with the reciprocal reinforced/non-reinforced odor combination. Odors used for conditioning were 3-octanol (8 μl in 10 ml mineral oil), 4-methylcyclohexanol (9 μl in 8 ml mineral oil) and isoamyl acetate (16 μl in 10 ml mineral oil). Only 3-octanol and 4-methylcyclohexanol were used during testing.

### Immunohistochemistry and confocal microscopy

4-6 day old female fly brains were dissected in ice-cold PBS and fixed in 4% paraformaldehyde solution in PBS at room temperature for 20 min. Fixed brains were then incubated in a blocking solution in 10% donkey serum (Jackson Immunoresearch) in PBS-triton (PBST) 0.5% for 30 min at room temperature. Brains were then incubated with primary antibodies, α-Bruchpilot, nc82, (1:30, mouse, DSHB) and α-GFP (1:1000, Chicken, ABCAM) in PBST for 72 h at 4°C. After 3 x 10 min washes in PBST, brains were incubated with secondary antibodies, Alexa Fluor 647-conjugated donkey α-mouse and Alexa Fluor 488-conjugated donkey α-chicken (both at 1:400; Jackson Immunoresearch) overnight at 4°C. After 3 x 10 min washes in PBS, brains were mounted in Vectashield (Vector Labs) on a glass slide. Fly brains were imaged with a Leica SP8-UV confocal microscope. Image resolution was 1024 x 512 with 0.5 μm step size and a frame average of 3 with a 40x objective. All images were analyzed using Fiji ^99^.

### *In vivo* two-photon calcium imaging

Functional-imaging experiments were performed as described previously ^44,50,59^ with some minor modifications. 1-day old flies were transferred into vials containing standard food (maximum of 30 flies per vial) and imaged 4-7 days later. Flies were briefly immobilized on ice (30-60 s) and mounted in a custom-made chamber allowing free antennae and leg movement. The head capsule was opened under room temperature buffer solution (5 mM TES, 103 mM NaCl, 3 mM KCl, 1.5 mM CaCl_2_, 4 mM MgCl_2_, 26 mM NaHCO_3_, 1 mM NaH_2_PO_4_, 8 mM Trehalose, 10mM glucose, pH 7). Membranes and trachea above the recording areas were manually removed. Individual flies in the recording chamber were placed under a two-photon microscope (Zeiss LSM 710mp) and an electric shock grid (in copper) was positioned in contact with the fly’s legs (visualized with a camera AV MAKO U-029B (Stemmer Imaging)). For all flies, GCaMP6m fluorescence signal was measured in a randomly chosen brain hemisphere. Fluorescence was excited by a Ti-Sapphire laser (Chameleon Ultra II, Coherent) using ∼140fs pulses, 80MHz repetition rate and centered at 920 nm. Images of 256 x 256 pixels were acquired at 6.34Hz controlled with Zen software (Zeiss). Odors were delivered to the fly on a clean air stream at 0.9mL/min using the 206A olfactometer (Aurora Scientific). Electric shocks were delivered with a DS2A isolated voltage stimulator controlled with a DG2A train/delay generator (both from Digitimer, Ltd). Imaging (Zen software), electric shocks and odor delivery were all controlled via TTL signals using an Arduino board (Arduino uno Rev3, Arduino.cc) combined with a project board (K&H-102) and Arduino custom-made codes.

For imaging electric shock responses in DANs, each fly was recorded for 30 s before the onset of a 1 min sequence of 12 x 1.5 s electric shocks of a given voltage at 0.2 Hz. Note that each fly was only tested for their response to one voltage intensity.

For imaging odor-evoked responses in MBONs following a differential training protocol, flies were first trained under the microscope with the same protocol as in Fig. 1a and Fig. 2b.

Flies were exposed to odor X for 1 min paired with 12 x 1.5 s electric shocks at 0.2 Hz (CS+), followed by 45 s of clean air and a second odor Y unpaired (CS-) for 1 min. Recording during the test started 1 min after the conditioning protocol. Flies were then exposed twice to a 5 s presentation of the CS+ and CS- with 30 s clean air exposure in between every odor (see **Fig. 2b** and **Supplementary Fig. 2**). Note that each fly was trained and recorded only once with a randomly chosen intensity of electric shock voltage during conditioning.

For imaging odor-evoked responses in PAM-β’2aγ5n DANs (MB109B-GAL4), MBON-γ2α’1 (MB077B-GAL4), MBON-γ1ped>αβ (MB112C-GAL4) and MBON-γ5β’2a (R66C08-GAL4) during training, flies were subjected to the following protocol (same as for the behavioral experiments in Fig. 3, 4 and 5): 1 min odor X unpaired (CS-) followed by 45 s of clean air, then 1 min presentation of second odor Y paired with 12 x 1.5 s 60V electric shocks at 0.2 Hz, followed by 45 s of clean air and 1 min of a third odor Z paired with 12 x 1.5 s 30V electric shocks at 0.2 Hz. Note that each fly was trained and recorded only once. Recorded images were manually segmented with Fiji using a custom-made code including an image stabilizer plugin ^100^. For each recording, one region of interest (ROI) was drawn around the zone expressing GCaMP6m (dendrites for MBON-γ1ped and MBON-γ2α’1 or axonal terminals for all DANs and for MBON-γ5β’2a) after image stabilization to generate the summed fluorescence at each frame. A second ROI of the same size was chosen in the background where no changes occur during the whole recording. The GCaMP6m fluorescence (F(t)) was then calculated by subtracting the background. Flies that did not respond to any odor presentations were excluded from the analysis. For subsequent analyses custom MATLAB (MathWorks, Inc) scripts were used to first calculate variation of calcium transient from the baseline F0 with the following equation: ΔF = (F(t) – F0) / F0. For DANs recordings, the baseline fluorescence, (F0), was defined as the mean fluorescence (F) from the 25 s before the start of the electric shock sequence. We chose this long duration for F0 as PPL1 DANs display rhythmic slow oscillatory activity ^101^ that could affect our ΔF calculation. For MBON recordings during training and testing, or PAM-β’2aγ5n DANs during training, F0 was defined as the mean fluorescence of the 2 s before each odor presentation. As we presented the CS+ and CS- twice in Fig. 2 and supplementary Fig. 2 to record MBON odor-evoked responses after learning, we used MATLAB to average the odor responses of the two odor presentations. In all experiments we calculated the mean fluorescence and the area under the curve (integral F/F0) during the whole sequence of electric shocks or odor presentation.

## QUANTIFICATION AND STATISTICAL ANALYSIS

All statistical analyses were performed using PRISM 9.1.2 (GraphPad Software). All behavioral and imaging data were first tested for normality using the D’Agostino and Pearson normality test. Normally distributed data were analyzed with parametric one-way ANOVA followed by Tukey’s honest significant difference (HSD) post hoc test. For non-Gaussian distributed data, Kruskal-Wallis test was performed followed by Dunn’s multiple comparisons test. Simple linear regression was used to fit a model to our data (shock responses or CS+ CS- difference). We also compared linear regression slopes with Prism (equivalent method to an Analysis of covariance). For the correlation between calcium transient and shock intensities we used parametric Pearson or non-parametric Spearman correlation. Paired t- or Wilcoxon tests were used to compare MBON CS+ and CS- responses after training. Unpaired t- or Mann-Whitney tests were used to compare cell body counting and quarters of area under curve for fluorescence data during training. All graphics were generated with Inkscape 1.0.2 (Inkscape). All statistical comparisons are presented in **Table S2**.

## AUTHOR CONTRIBUTIONS

E.P. and S.W. conceived the project and M.V., M.P.D. and E.P. designed all experiments. E.P., M.V., N.M. and S.P. performed behavioral experiments. E. P. and M.V. analyzed all behavioral experiments. M.P.D., E.P., A.P., P.J. and M.A. performed imaging experiments. E.P., M.A. and M.P.D. analyzed imaging data. Anatomical data were collected by M.V. and E.P. The manuscript was written by E.P., M.V., M.P.D. and S.W.

## Acknowledgments

We thank Stephanie Trouche, Johannes Felsenberg and Suewei Lin for comments on the manuscript. We thank the Bloomington stock center for flies. We thank Yaling Huang for making the R56H09-GAL80 construct. We thank Richard Griffith and Luis D. Suarez for help with behavioral experiments. E.P. is funded by the ATIP-Avenir program from CNRS and Inserm, the Bettencourt Schueller Foundation, the French Research Agency (ANR-PRC 2021) and the Institute of Functional Genomics, Montpellier, France. S.W. is funded by a Wellcome Trust Principal Research Fellowship in the Basic Biomedical Sciences (200846/Z/16/Z) and a Wellcome Trust Collaborative Award (2023261/Z/16/Z). We thank the Montpellier Biocampus facilities (University of Montpellier, CNRS, INSERM, Montpellier, France) MRI, Drosophila facility and IPAM for the confocal microscope, the fly food and the 2-photon microscope, respectively.

---

**Supplementary Fig. 1.**
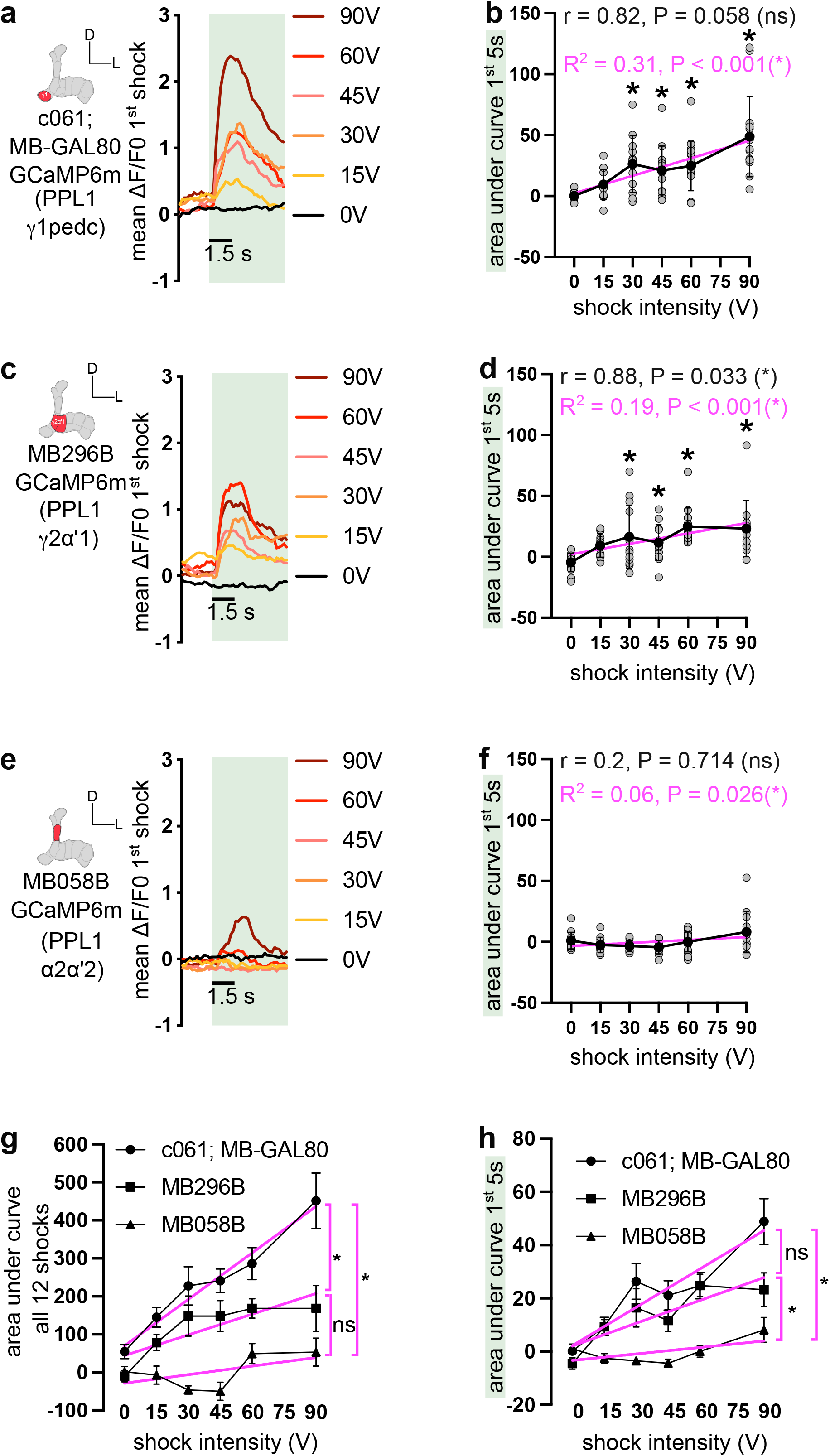
Further analysis of PPL1-γ1ped, PPL1-γ2α’1 and PPL1-α2α’2 DAN responses to electric shocks of different intensity (related to Fig. 1). **a, c and e.** Mean ΔF/F0 calcium traces during the first 5 s (light green rectangle) for the 1^st^ shock presentation of 0V, 15V, 30V, 45V, 60V or 90V for PPL1-γ1ped targeted with c061-GAL4;MBGAL80 (**a**), PPL1-γ2α’1 targeted with MB296B-GAL4 (**c**), and PPL1-α2α’2 targeted with MB058B-GAL4 (**e**). **b.** Mean area under the curve ± SD during the 1^st^ 5 s of PPL1-γ1ped DAN responses showing a correlation (Spearman correlation, close to significance) with the shock intensity. **d.** Mean area under the curve ± SD during the 1^st^ 5s of PPL1-γ2α’1 DAN responses showing a significant correlation (Spearman correlation) with the shock intensity. **f.** Mean area under the curve ± SD during the 1^st^ 5s of PPL1-α2α’2 DAN responses showing no correlation (Spearman correlation) with the shock intensity. **g.** Mean area under the curve ± SEM during all 12 shocks of PPL1-γ1ped (c061;MBGAL80), PPL1-γ2α’1 (MB296B) and PPL1-α2α’2 (MB058B) responses with corresponding linear regression (*y* = 0.61*x* + 31.87, R^2^=0.35; *y* = 0.55*x* + 27.44, R^2^ = 0.12 and *y* = 0.33*x* + 16.81, R^2^=0.06, respectively). **h.** Mean area under the curve ± SEM during the 1^st^ 5s of PPL1-γ1ped (c061;MBGAL80), PPL1-γ2α’1 (MB296B) and PPL1-α2α’2 (MB058B) responses with corresponding linear regression (*y* = 0.08*x* + 4.37, R^2^ = 0.29; *y* = 0.07*x*+ 3.68, R^2^ = 0.18 and *y* = 0.03*x* + 1.91, R^2^ = 0.05, respectively). Slopes are compared to zero with the F test. N for each group are provided in Fig. 1. Individual data points are displayed as dots. *P<0.05. ns: non-significant.

**Supplementary Fig. 2.**
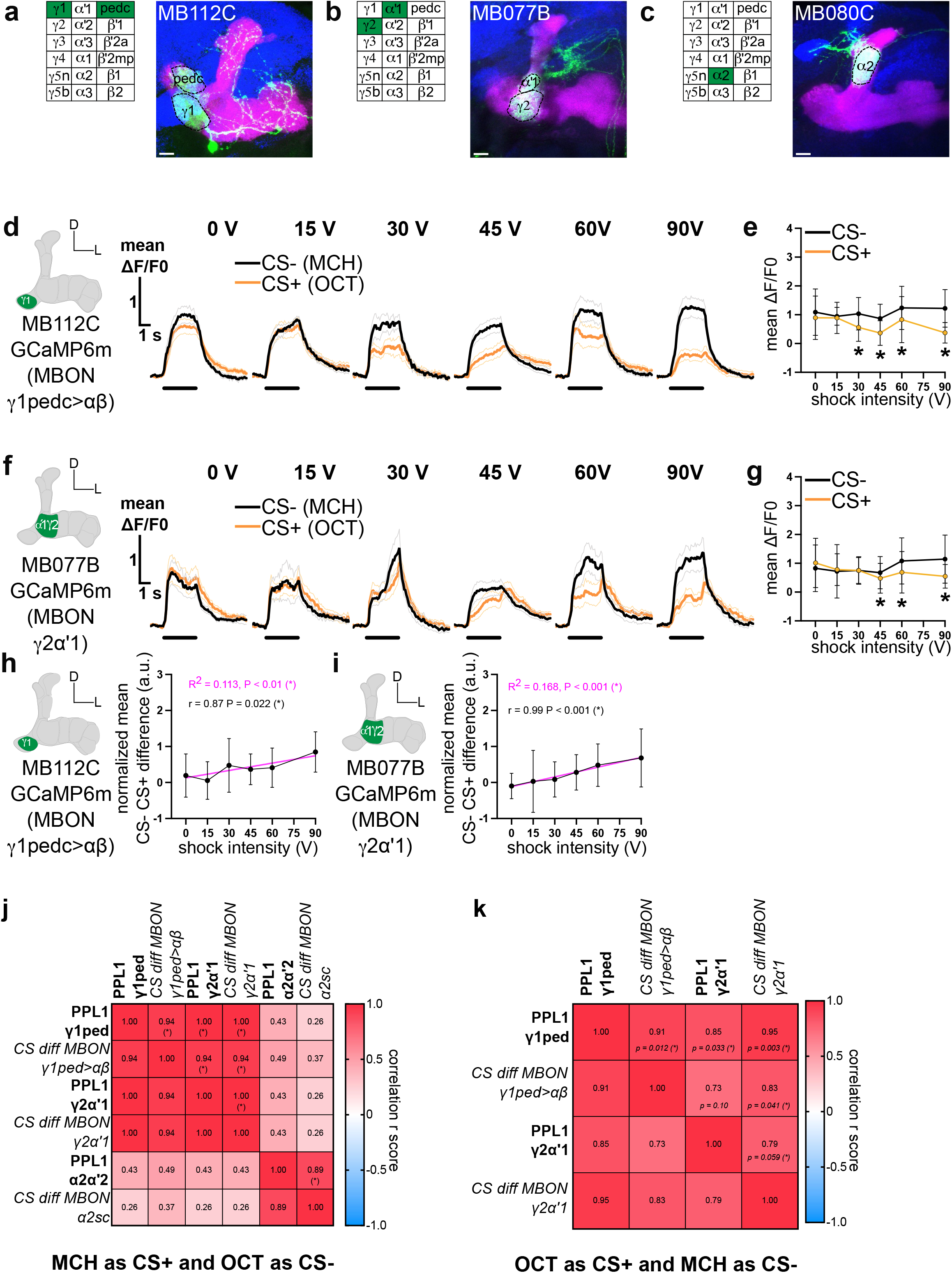
Aversive learning drives intensity-dependent plasticity of MBON-γ1ped >αβ and MBON-γ2α’1 odor responses, using the reciprocal odor combination (related to Fig. 2). **a-c.** Z-projection of confocal stacks from fly brains harboring (**a**) MB112C-GAL4-, (**b**) MB077B-GAL4 and (**c**) MB080C-GAL4-driving UAS-mCD8::GFP labelling one MBON per hemisphere innervating specific MB compartments, and summarized with an inset. MB co-labelled with 247-LexA::VP16-driven lexAop-rCD2::mRFP (magenta) and the whole brain with the presynaptic marker anti-nc82/anti-Bruchpilot (blue). Scale bar 10μm. **d and f.** Mean ΔF/F0 calcium transients ± SEM for the CS- (average of the two presentations, black) and for the CS+ (average of the two presentations, orange) immediately after training the CS+ with either 0V, 15V, 30V, 45V, 60V or 90V. MB112C is used to target MBON-γ1ped >αβ (**d**) and MB077B for MBON-γ2α’1 (**f**). **e and g.** Mean ΔF/F0 ± SD during the 5 s odor presentation for the CS- (black) and for the CS+ (orange) when paired with either 0V, 15V, 30V, 45V, 60V or 90V for MBON-γ1ped >αβ(N= 26, 15, 16, 14, 15 and 14) (**e**) and MBON-γ2α’1 (N= 18, 16, 15, 15, 15 and 16) (**g**). Statistical tests are unpaired t-test (or Wilcoxon test) between the CS- and the CS+ for each voltage intensity. **h and i.** Normalized CS- CS+ difference ± SD and fitted linear regression (magenta) for MBON-γ1ped >αβ(**h**) and MBON-γ2α’1 (**i**) with a Pearson (r) (or Spearman) correlation. *P<0.05. ns: non-significant. **j.** Correlation matrix for all PPL1-γ1ped, PPL1-γ2α’1 and PPL1-α2α’2 DAN shock responses with MBON-γ1ped >αβ, MBON-γ2α’1 and MBON-α2sc CS- CS+ difference odor-evoked responses following aversive learning (MCH as CS+ and OCT as CS-). PPL1 DAN shock responses are strongly correlated (Spearman test) with MBONs odor-evoked responses in the γ1ped and γ2α’1 MB compartments but not in the α2 compartment. **k.** Correlation matrix for both PPL1-γ1ped and PPL1-γ2α’1 DAN shock responses with the MBON-γ1ped >αβ and MBON-γ2α’1 CS- CS+ difference in odor-evoked responses following aversive learning (OCT as CS+ and MCH as CS-). PPL1-γ1ped and PPL1-γ2α’1 DAN shock responses are strongly correlated (Pearson test) with MBON odor-evoked responses in the γ1ped and γ2α’1 MB compartments. *P<0.05.

**Supplementary Fig. 3.**
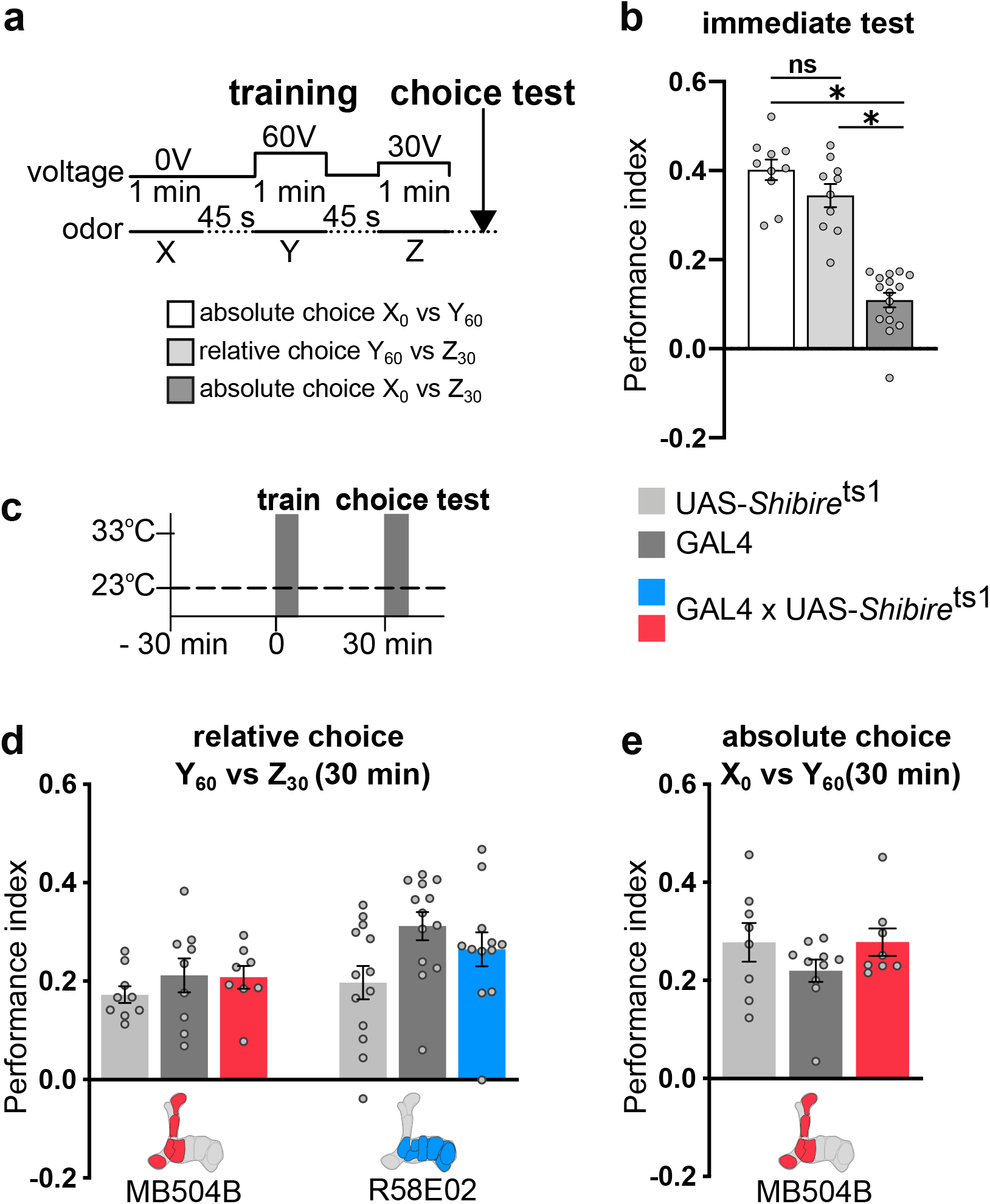
Control experiments related to Fig. 3. **a.** Training protocol. **b.** No difference in immediate memory performance of control wild-type Canton-S flies for relative choice (Y_60_ vs Z_30_, in light gray) vs absolute choice (X_0_ vs Y_60_, in white) but vs absolute choice (X_0_ vs Z_30_, in dark gray) (Kruskal-Wallis, N=10-16, P<0.05). **c.** Training protocol and control permissive temperature procedure for *Shi*^ts1^ manipulations. **d-e.** No statistical differences were apparent between any groups and their relevant controls for MB504B-GAL4 or R58E02-GAL4 during relative (**d**) or absolute (**e**) choices (MB504B-GAL4: 1-way ANOVA, N=8-9, P>0.05 and Kruskal-Wallis, N=8-10, P=0.049 (but no post-hoc difference between the groups) respectively; R58E02-GAL4: 1-way ANOVA, N=12-13, P>0.05). Data are mean ± SEM. Individual data points are displayed as dots. *P<0.05, ns: non-significant.

**Supplementary Fig. 4.**
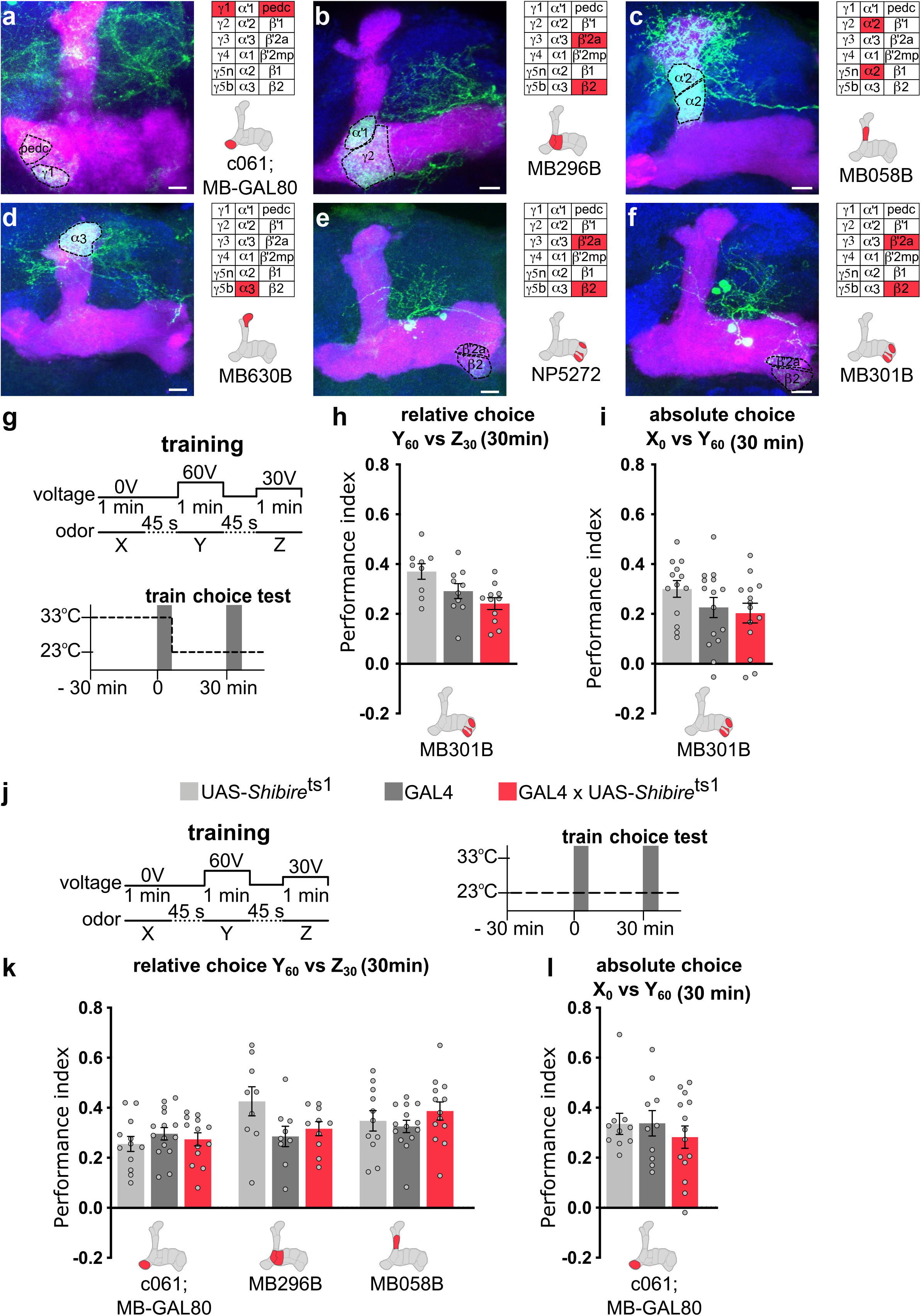
GAL4 line expression patterns and control experiments related to Fig. 4. **a-f.** Z-projection of (**a**) c061-GAL4;MBGAL80 (**b**) MB296B-GAL4, (**c**) MB058B-GAL4, (**d**) MB630B-GAL4, (**e**) NP5272-GAL4 and (**f**) MB301B-GAL4 driving UAS-mCD8::GFP labelling a single type of punishment DAN (schematized in red with the MB in gray and summarized with an inset). MB co-labelled with 247-LexA::VP16-driven lexAop-rCD2::mRFP (magenta) and the whole brain with the presynaptic marker anti-Bruchpilot, nc82 (blue). Scale bar 10μm. **g.** Training protocol and temperature shifting procedure for *Shi*^ts1^ manipulations. **h and i.** Blocking output from MB301B-GAL4 (PAM-β2β’2a) during training does not alter 30 min relative (**h**) or absolute (**i**) choice compared to that of relevant controls (1-way ANOVA, N=9-11, P<0.05; 1-way ANOVA, N=13-15, P>0.05; respectively). **j.** Training protocol and control permissive temperature procedure for *Shi*^ts1^ manipulations. **k and l.** No statistical differences were apparent between any groups and their relevant controls for c061;MBGAL80, MB296B-GAL4 or MB058B-GAL4 during relative (**k**) or absolute (**l**) choices (c061;MBGAL80: 1-way ANOVA, N=12-15, P>0.05 (relative) and Kruskal-Wallis, N=10-14, P>0.05 (absolute); MB296B: 1-way ANOVA, N=9-10, P>0.05; MB058B: Kruskal-Wallis, N=11-14, P>0.05). Data are mean ± SEM. Individual data points are displayed as dots.

**Supplementary Fig. 5.**
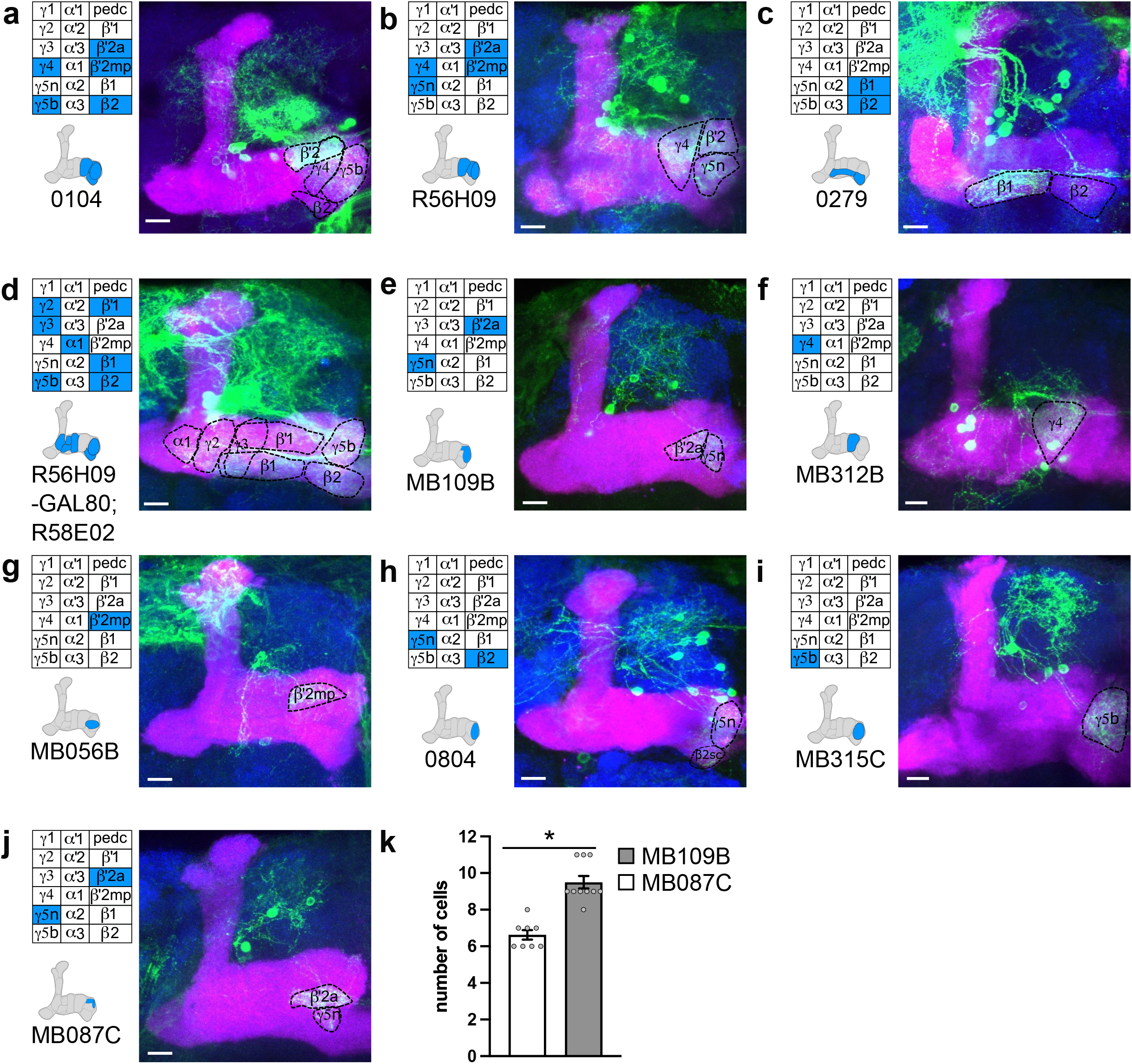
GAL4 line expression patterns related to Fig. 5. **a-j.** Z-projection of (**a**) 0104-GAL4, (**b**) R56H09-GAL4, (**c**) 0279-GAL4, (**d**) R56H09-GAL80;R58E02-GAL4 (**e**) MB109B-GAL4, (**f**) MB312B-GAL4, (**g**) MB056B-GAL4, (**h**) 0804-GAL4, (**i**) MB315C-GAL4 and (**j**) MB087C-GAL4 driving UAS-mCD8::GFP labelling a combination, or a single type, of PAM DANs per hemisphere, innervating specific MB compartments (schematized in blue with the MB in gray and summarized with an inset). MB co-labelled with 247-LexA::VP16-driven lexAop-rCD2::mRFP (magenta) and the whole brain with the presynaptic marker anti-Bruchpilot, nc82 (blue). Scale bar 10μm. **k.** GFP positive cell body counting showing a significant higher number of cell bodies for MB109B-GAL4 compared to that of MB087C-GAL4 (Mann-Whitney test, P<0.05). Data are mean ± SEM. Individual data points (left or right hemisphere combined) are displayed as dots. *P<0.05.

**Supplementary Fig. 6.**
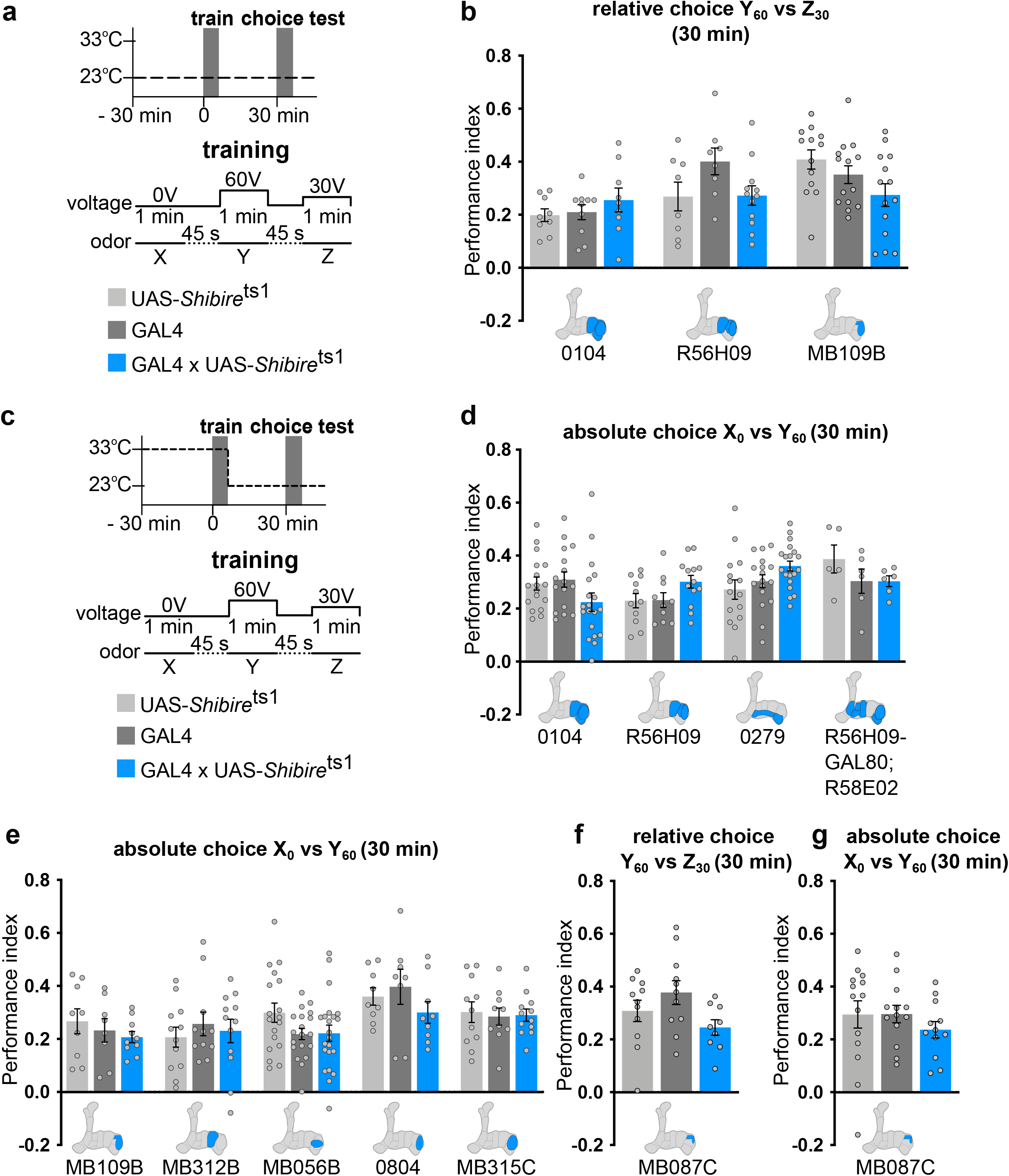
UAS-*Shi*^ts1^ control experiments related to Fig. 5. **a.** Training protocol and control permissive temperature procedure for *Shi*^ts1^ manipulations. **b.** No statistical differences were apparent between any groups and their relevant controls for 0104, R56H09-GAL4 or MB109B-GAL4 during relative choice (1-way ANOVA, N=9-10, P>0.05, 1-way ANOVA, N=8-12, P>0.05 and 1-way ANOVA, N=13-14, P>0.05; respectively). **c.** Training protocol and temperature shifting procedure for *Shi*^ts1^ manipulations (for **d-g**). **d.** Blocking output from 0104-GAL4, R56H09-GAL4, 0279-GAL4 or R56H09-GAL80;R58E02-GAL4 during training did not impair 30 min absolute choice compared to that of relevant controls (Kruskal-Wallis, N=16-19, P>0.05; 1-way ANOVA, N=10-13, P>0.05; 1-way ANOVA, N=15-20, P>0.05; Kruskal-Wallis, N=5-6, P>0.05; respectively). **e.** Blocking output from MB109B-GAL4, MB312B-GAL4, MB056B-GAL4, 0804-GAL4 or MB315C-GAL4 during training did not impair 30 min absolute choice compared to that of relevant controls (1-way ANOVA, N=9-10, P>0.05; 1-way ANOVA, N=11-12, P>0.05; 1-way ANOVA, N=18-22, P>0.05; 1-way ANOVA, N=8-9, P>0.05; 1-way ANOVA, N=10-12, P>0.05, respectively). **f-g.** Blocking output from MB087C-GAL4 during training did not impair 30 min relative (**f**) or absolute (**g**) choices compared to that of relevant controls (1-way ANOVA, N=9-11, P>0.05; 1-way ANOVA, N=12-14, P>0.05, respectively). Data are mean ± SEM. Individual data points are displayed as dots.

**Supplementary Fig. 7.**
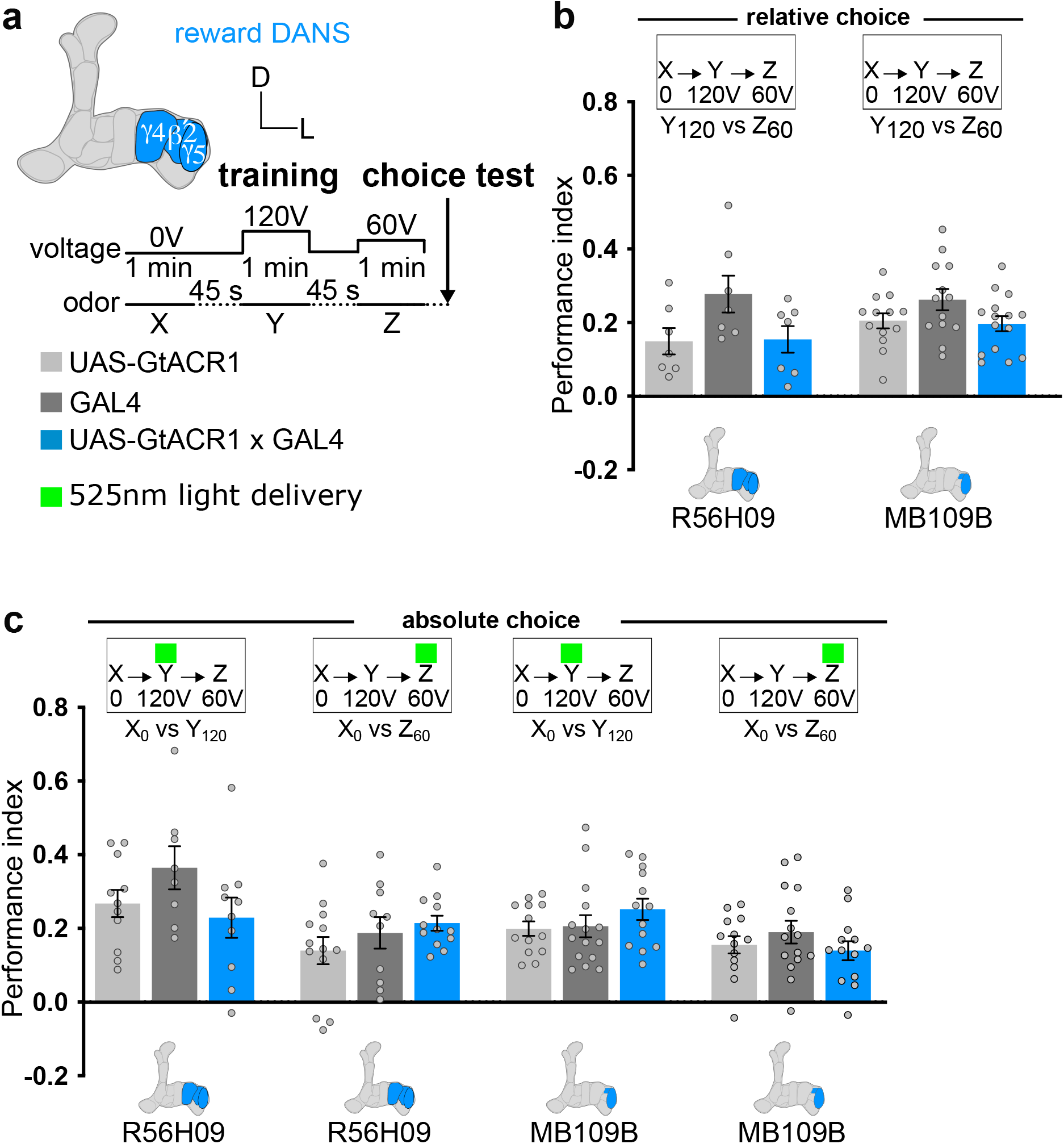
GtACR1 control experiments related to Fig. 5. **a.** Schematics of relevant reward DAN innervation for R56H09- and MB109B-GAL4 (schematized in blue with the MB in gray) and training protocol. **b.** No delivery of 525nm green light during training did not impair relative choice (Y_120_ vs Z_60_) for both R56H09- and MB109B-GAL4 lines expressing UAS-GtACR1 (Kruskal-Wallis test, N=7; P>0.05; 1-way ANOVA, N=13-15, P>0.05, respectively). **c.** Delivering 525nm green light during training Y + 120V or Z + 60V had no effect on absolute X_0_ vs Y_120_ or X_0_ vs Z_60_ immediate relative choices for both R56H09- and MB109B-GAL4 lines expressing UAS-GtACR1 (1-way ANOVA, N=8-11, P>0.05; 1-way ANOVA, N=10-13, P>0.05; 1-way ANOVA, N=13-15, P>0.05; 1-way ANOVA, N=13-15, P>0.05, respectively). Data are mean ± SEM. Individual data points are displayed as dots.

**Supplementary Fig. 8.**
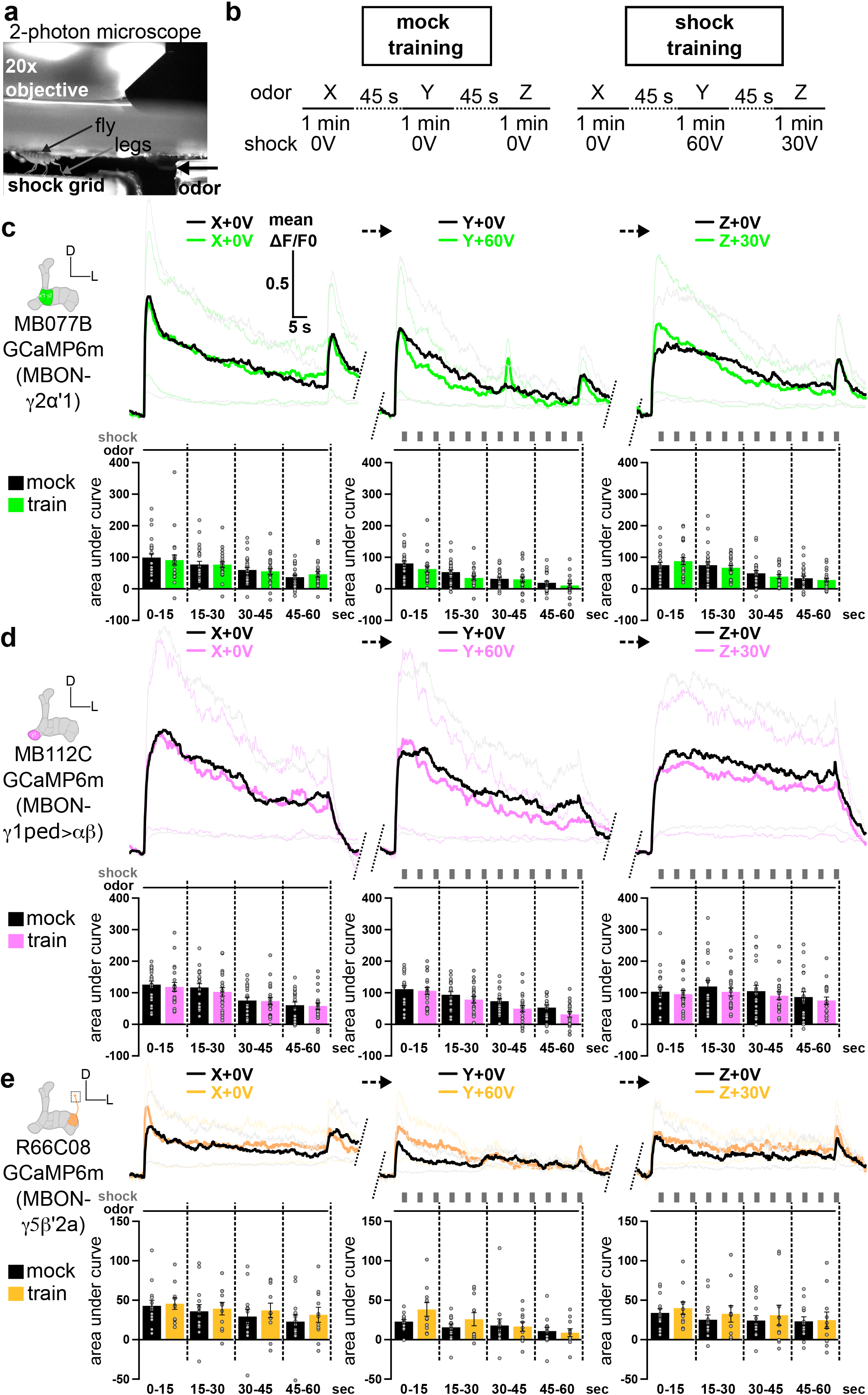
MBON-γ2α’1, MBON-γ1ped>αβ and MBON-γ5β’2a responses during relative training (related to Fig. 6). **a.** Experimental setup. **b.** Control mock protocol: the odors X, Y and Z are presented alone. For shock training: the odor X is unpaired, odor Y is paired with 60V, and odor Z with 30V. **c-e.** GCaMP6m calcium traces for MBON-γ2α’1 (**c**, N=26 mock and N=23 train), MBON-γ1ped>αβ(**d**, N=21 mock and N=21 train) and MBON-γ5β’2a (**e**, N=14 mock and N=11 train). Mean area under curve quantifications for each 15 sec segment of every 1 min odor presentation for MBON-γ2α’1, MBON-γ1ped>αβand MBON-γ5β’2a show no significant differences between shock training compared to that of mock control training (All unpaired t or Mann-Whitney test, P > 0.05). Data are mean ± SEM. Individual data points are displayed as dots.

**Supplementary Fig. 9.**
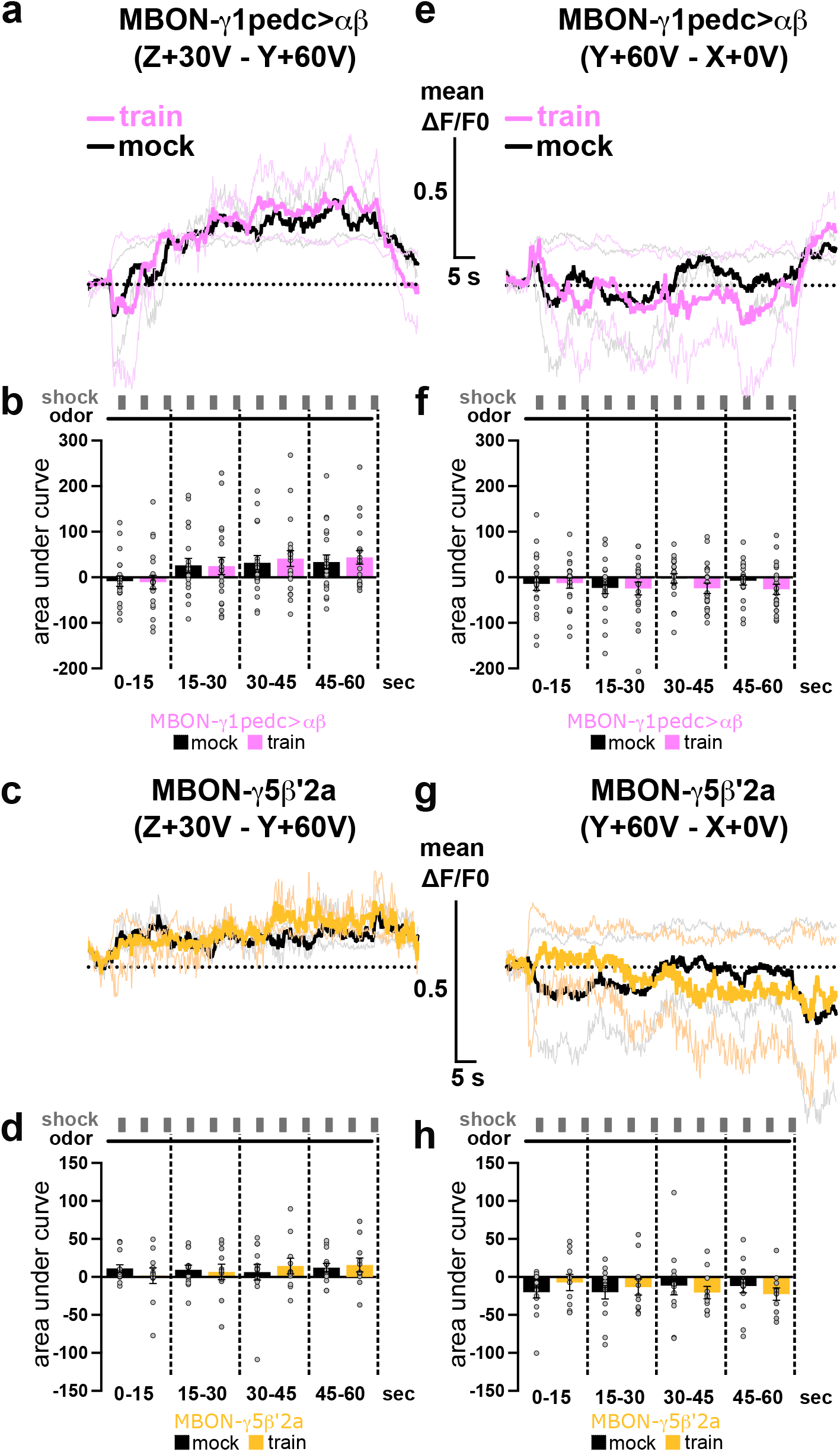
MBON-γ1ped>αβ and MBON-γ5β’2a differential responses during training (related to Fig. 6). **a-b.** Mean subtracted Z+30V – Y+60V calcium traces and mean area under curve for MBON-γ1ped>αβduring mock (N=21) or shock training (N=21) protocols. No significant differences were found for any quarter of the Z+30V – Y+60V odor presentation subtraction in the trained group compared to that of the mock group (All unpaired t or Mann-Whitney tests; P>0.05). **c-d.** Mean subtracted Z+30V – Y+60V calcium traces and mean area under the curve for MBON-γ5β’2a during mock (N=14) or shock training (N=11) protocols. No significant differences were found for any quarter of the Z+30V – Y+60V odor presentation subtraction in the trained group compared to that of the mock group (All unpaired t or Mann-Whitney tests; P>0.05). **e-f.** Mean subtracted Y+60V – X+0V calcium traces and mean area under the curve for MBON-γ1ped>αβ during mock (N=21) or shock training (N=21) protocols. No significant differences were found for any quarter of the Y+60V – X+0V odor presentation subtraction in the trained group compared to that of the mock group (All unpaired t or Mann-Whitney tests; P>0.05). **g-h.** Mean subtracted Y+60V – X+0V calcium traces and mean area under the curve for MBON-γ5β’2a during mock (N=14) or shock training (N=11) protocols. No significant differences were found for any quarter of the Y+60V – X+0V odor presentation subtraction in the train group compared to that of the mock group (All unpaired t or Mann-Whitney tests; P>0.05). Data are mean ± SEM. Individual data points are displayed as dots.

## References

1. Rangel, A., Camerer, C. & Montague, P. R. A framework for studying the neurobiology of value-based decision making. Nat. Rev. Neurosci. 9, 545–556 (2008).

2. Pavlov, I. P. Conditioned Reflexes. (1927).

3. Tremblay, L. & Schultz, W. Relative reward preference in primate orbitofrontal cortex. Nature 398, 704–708 (1999).

4. Seymour, B. & McClure, S. M. Anchors, scales and the relative coding of value in the brain. Curr. Opin. Neurobiol. 18, 173–178 (2008).

5. Hunter, L. E. & Daw, N. D. Context-sensitive valuation and learning. Curr. Opin. Behav. Sci. 41, 122–127 (2021).

6. Tobler, P. N., Fiorillo, C. D. & Schultz, W. Adaptive coding of reward value by dopamine neurons. Science 307, 1642–5 (2005).

7. Diederen, K. M. J. et al. Dopamine modulates adaptive prediction error coding in the human midbrain and striatum. J. Neurosci. 37, 1708–1720 (2017).

8. Padoa-Schioppa, C. & Assad, J. A. Neurons in the orbitofrontal cortex encode economic value. Nature 441, 223–226 (2006).

9. Lim, S. L., O’Doherty, J. P. & Rangel, A. The decision value computations in the vmPFC and striatum use a relative value code that is guided by visual attention. J. Neurosci. 31, 13214–13223 (2011).

10. Strait, C. E., Sleezer, B. J. & Hayden, B. Y. Signatures of Value Comparison in Ventral Striatum Neurons. PLOS Biol. 13, e1002173 (2015).

11. Hebscher, M., Barkan-Abramski, M., Goldsmith, M., Aharon-Peretz, J. & Gilboa, A. Memory, Decision-Making, and the Ventromedial Prefrontal Cortex (vmPFC): The Roles of Subcallosal and Posterior Orbitofrontal Cortices in Monitoring and Control Processes. Cereb. Cortex bhv220 (2015). doi:10.1093/cercor/bhv220

12. Saez, R. A., Saez, A., Paton, J. J., Lau, B. & Salzman, C. D. Distinct Roles for the Amygdala and Orbitofrontal Cortex in Representing the Relative Amount of Expected Reward. Neuron 95, 70–77.e3 (2017).

13. Lak, A., Stauffer, W. R. & Schultz, W. Dopamine neurons learn relative chosen value from probabilistic rewards. Elife 5, 1–19 (2016).

14. Klein, T. A., Ullsperger, M. & Jocham, G. Learning relative values in the striatum induces violations of normative decision making. Nat. Commun. 8, 1–12 (2017).

15. Padoa-Schioppa, C. & Conen, K. E. Orbitofrontal Cortex: A Neural Circuit for Economic Decisions. Neuron 96, 736–754 (2017).

16. Brooks, A. M. et al. From bad to worse: Striatal coding of the relative value of painful decisions. Front. Neurosci. 4, 1–8 (2010).

17. Perisse, E. et al. Different Kenyon Cell Populations Drive Learned Approach and Avoidance in Drosophila. Neuron 79, 945–956 (2013).

18. Hosokawa, T., Kato, K., Inoue, M. & Mikami, A. Neurons in the macaque orbitofrontal cortex code relative preference of both rewarding and aversive outcomes. Neurosci. Res. 57, 434–445 (2007).

19. Campese, V. D. et al. Chemogenetic inhibition reveals that processing relative but not absolute threat requires basal amygdala. J. Neurosci. 39, 8510–8516 (2019).

20. Rescorla, R. a & Wagner, a R. A theory of Pavlovian conditioning: Variations in the effectiveness of reinforcement and nonreinforcement. Class. Cond. II Curr. Res. Theory 21, 64–99 (1972).

21. Sutton, R. S. & Barto, A. G. Reinforcement Learning : An Introduction. (1998).

22. Schultz, W. A Neural Substrate of Prediction and Reward. Science (80-. ). 275, 1593–1599 (1997).

23. Glimcher, P. W. Understanding dopamine and reinforcement learning: The dopamine reward prediction error hypothesis. Proc. Natl. Acad. Sci. 108, 17647–15654 (2011).

24. Watabe-Uchida, M., Eshel, N. & Uchida, N. Neural Circuitry of Reward Prediction Error. Annu. Rev. Neurosci. 40, 373–394 (2017).

25. Waddell, S. Reinforcement signalling in Drosophila ; dopamine does it all after all. Curr. Opin. Neurobiol. 23, 324–329 (2013).

26. Watabe-Uchida, M. & Uchida, N. Multiple dopamine systems: Weal and woe of dopamine. Cold Spring Harb. Symp. Quant. Biol. 83, 83–95 (2018).

27. Brooks, A. M. & Berns, G. S. Aversive stimuli and loss in the mesocorticolimbic dopamine system. Trends Cogn. Sci. 17, 281–286 (2013).

28. Adel, M. & Griffith, L. C. The Role of Dopamine in Associative Learning in Drosophila: An Updated Unified Model. Neurosci. Bull. 37, 831–852 (2021).

29. Heisenberg, M. Mushroom body memoir: from maps to models. Nat. Rev. Neurosci. 4, 266–275 (2003).

30. Waddell, S. Dopamine reveals neural circuit mechanisms of fly memory. Trends Neurosci. 33, 457–464 (2010).

31. Aso, Y. et al. The neuronal architecture of the mushroom body provides a logic for associative learning. Elife 3, 1–47 (2014).

32. Aso, Y. et al. Mushroom body output neurons encode valence and guide memory-based action selection in Drosophila. Elife 3, e04580 (2014).

33. Schwaerzel, M. et al. Dopamine and octopamine differentiate between aversive and appetitive olfactory memories in Drosophila. J. Neurosci. 23, 10495–502 (2003).

34. Claridge-Chang, A. et al. Writing Memories with Light-Addressable Reinforcement Circuitry. Cell 139, 405– 415 (2009).

35. Aso, Y. et al. Specific Dopaminergic Neurons for the Formation of Labile Aversive Memory. Curr. Biol. 20, 1445–1451 (2010).

36. Aso, Y. et al. Three Dopamine Pathways Induce Aversive Odor Memories with Different Stability. PLoS Genet. 8, e1002768 (2012).

37. Aso, Y. & Rubin, G. M. Dopaminergic neurons write and update memories with cell-type-specific rules. Elife 5, 1–15 (2016).

38. Takemura, S. ya, et al. A connectome of a learning and memory center in the adult Drosophila brain. Elife 6, 1–43 (2017).

39. Liu, C. et al. A subset of dopamine neurons signals reward for odour memory in Drosophila. Nature 488, 512– 516 (2012).

40. Burke, C. J. et al. Layered reward signalling through octopamine and dopamine in Drosophila. Nature 492, 433–437 (2012).

41. Huetteroth, W. et al. Sweet Taste and Nutrient Value Subdivide Rewarding Dopaminergic Neurons in Drosophila. Curr. Biol. 25, 751–758 (2015).

42. Yamagata, N. et al. Distinct dopamine neurons mediate reward signals for short- and long-term memories. Proc. Natl. Acad. Sci. 112, 578–583 (2015).

43. Lin, S. et al. Neural correlates of water reward in thirsty Drosophila. Nat. Neurosci. 17, 1536–1542 (2014).

44. Felsenberg, J. et al. Integration of Parallel Opposing Memories Underlies Memory Extinction. Cell 175, 709–722.e15 (2018).

45. Jacob, P. F. & Waddell, S. Spaced Training Forms Complementary Long-Term Memories of Opposite Valence in Drosophila. Neuron (2020). doi:10.1016/j.neuron.2020.03.013

46. McCurdy, L. Y., Sareen, P., Davoudian, P. A. & Nitabach, M. N. Dopaminergic mechanism underlying reward-encoding of punishment omission during reversal learning in Drosophila. Nat. Commun. 12, (2021).

47. Wang, Y., Mamiya, A., Chiang, A. S. & Zhong, Y. Imaging of an early memory trace in the Drosophila mushroom body. J. Neurosci. 28, 4368–4376 (2008).

48. Honegger, K. S., Campbell, R. A. A. a. & Turner, G. C. Cellular-resolution population imaging reveals robust sparse coding in the drosophila mushroom body. J. Neurosci. 31, 11772–11785 (2011).

49. Tanaka, N. K., Tanimoto, H. & Ito, K. Neuronal assemblies of theDrosophila mushroom body. J. Comp. Neurol. 508, 711–755 (2008).

50. Perisse, E. et al. Aversive Learning and Appetitive Motivation Toggle Feed-Forward Inhibition in the Drosophila Mushroom Body. Neuron 90, 1086–1099 (2016).

51. Hattori, D. et al. Representations of Novelty and Familiarity in a Mushroom Body Compartment. Cell 169, 956–969.e17 (2017).

52. Cohn, R., Morantte, I. & Ruta, V. Coordinated and Compartmentalized Neuromodulation Shapes Sensory Processing in Drosophila. Cell 163, 1742–1755 (2015).

53. Li, F. et al. The connectome of the adult drosophila mushroom body provides insights into function. Elife 9, 1– 217 (2020).

54. Eschbach, C. et al. Circuits for integrating learned and innate valences in the insect brain. Elife 10, 1–36 (2021).

55. Eschbach, C. et al. Recurrent architecture for adaptive regulation of learning in the insect brain. Nat. Neurosci. (2020). doi:10.1038/s41593-020-0607-9

56. Séjourné, J. et al. Mushroom body efferent neurons responsible for aversive olfactory memory retrieval in Drosophila. Nat. Neurosci. 14, 903–910 (2011).

57. Boto, T., Louis, T., Jindachomthong, K., Jalink, K. & Tomchik, S. M. M. Dopaminergic Modulation of cAMP Drives Nonlinear Plasticity across the Drosophila Mushroom Body Lobes. Curr. Biol. 24, 822–831 (2014).

58. Hige, T., Aso, Y., Modi, M. N., Rubin, G. M. & Turner, G. C. Heterosynaptic Plasticity Underlies Aversive Olfactory Learning in Drosophila. Neuron 88, 985–998 (2015).

59. Owald, D. et al. Activity of Defined Mushroom Body Output Neurons Underlies Learned Olfactory Behavior in Drosophila. Neuron 86, 417–427 (2015).

60. Bouzaiane, E., Trannoy, S., Scheunemann, L., Plaçais, P.-Y. & Preat, T. Two Independent Mushroom Body Output Circuits Retrieve the Six Discrete Components of Drosophila Aversive Memory. Cell Rep. 1280–1292 (2015). doi:10.1016/j.celrep.2015.04.044

61. Berry, J. A., Phan, A. & Davis, R. L. Dopamine Neurons Mediate Learning and Forgetting through Bidirectional Modulation of a Memory Trace. Cell Rep. 25, 651–662.e5 (2018).

62. Handler, A. et al. Distinct Dopamine Receptor Pathways Underlie the Temporal Sensitivity of Associative Learning. Cell 178, 60–75.e19 (2019).

63. Cervantes-Sandoval, I., Davis, R. L. & Berry, J. A. Rac1 Impairs Forgetting-Induced Cellular Plasticity in Mushroom Body Output Neurons. Front. Cell. Neurosci. 14, 1–11 (2020).

64. Owald, D. & Waddell, S. Olfactory learning skews mushroom body output pathways to steer behavioral choice in Drosophila. Curr. Opin. Neurobiol. 35, 178–184 (2015).

65. Tully, T. & Quinn, W. G. Classical conditioning and retention in normal and mutant Drosophila melanogaster. (1985).

66. Yin, Y., Chen, N., Zhang, S. & Guo, A. Choice strategies in Drosophila are based on competition between olfactory memories. 30, 279–288 (2009).

67. Scheunemann, L. et al. AKAPS Act in a Two-Step Mechanism of Memory Acquisition. J. Neurosci. 33, 17422–17428 (2013).

68. Das, G. et al. Drosophila Learn Opposing Components of a Compound Food Stimulus. Curr. Biol. 24, 1723– 1730 (2014).

69. Galili, D. S. S. et al. Converging circuits mediate temperature and shock aversive olfactory conditioning in Drosophila. Curr. Biol. 24, 1712–1722 (2014).

70. Zhao, C. et al. Predictive olfactory learning in Drosophila. Sci. Rep. 11, (2021).

71. Mao, Z. & Davis, R. L. Eight different types of dopaminergic neurons innervate the Drosophila mushroom body neuropil: anatomical and physiological heterogeneity. Front. Neural Circuits 3, 5 (2009).

72. Dylla, K. V., Raiser, G., Galizia, C. G. & Szyszka, P. Trace conditioning in drosophila induces associative plasticity in mushroom body kenyon cells and dopaminergic neurons. Front. Neural Circuits 11, 1–14 (2017).

73. Brand, A. H. & Perrimon, N. Targeted gene expression as a means of altering cell fates and generating dominant phenotypes. Development 118, 401–15 (1993).

74. Chen, T.-W. et al. Ultrasensitive fluorescent proteins for imaging neuronal activity. Nature 499, 295–300 (2013).

75. Kitamoto, T. Conditional modification of behavior in drosophila by targeted expression of a temperature-sensitive shibire allele in defined neurons. J. Neurobiol. 47, 81–92 (2001).

76. Otto, N. et al. Input Connectivity Reveals Additional Heterogeneity of Dopaminergic Reinforcement in Drosophila. Curr. Biol. 30, 3200–3211.e8 (2020).

77. Vogt, K. et al. Shared mushroom body circuits underlie visual and olfactory memories in Drosophila. Elife 3, 3–5 (2014).

78. Mohammad, F. et al. Optogenetic inhibition of behavior with anion channelrhodopsins. Nat. Methods 14, 271– 274 (2017).

79. Weiglein, A. et al. Aversive teaching signals from individual dopamine neurons in larval Drosophila show qualitative differences in their temporal ‘fingerprint’. J. Comp. Neurol. (2020). doi:10.1002/cne.25037

80. Schroll, C. et al. Light-Induced Activation of Distinct Modulatory Neurons Triggers Appetitive or Aversive Learning in Drosophila Larvae. Curr. Biol. 16, 1741–1747 (2006).

81. Lewis, L. P. C. et al. A Higher Brain Circuit for Immediate Integration of Conflicting Sensory Information in Drosophila. Curr. Biol. 25, 2203–2214 (2015).

82. Ueoka, Y., Hiroi, M., Abe, T. & Tabata, T. Suppression of a single pair of mushroom body output neurons in Drosophila triggers aversive associations. FEBS Open Bio 7, 562–576 (2017).

83. Scaplen, K. M., Talay, M., Salamon, S., Nuñez, K. M. & Waterman, A. G. Circuits that encode and predict alcohol associated preference. 1–15 (2019).

84. Senapati, B. et al. A neural mechanism for deprivation state-specific expression of relevant memories in Drosophila. Nat. Neurosci. 22, 2029–2039 (2019).

85. Konorski, J. Integrative activity of the brain: An interdisciplinary approach. (1967). doi:10.1001/jama.1968.03140050055025

86. Dickinson, A. & Dearing, M. F. Appetitive-Aversive Interactions and Inhibitory Processes. Mechanisms of learning and motivation (1979).

87. Nasser, H. M. & McNally, G. P. Appetitive-aversive interactions in Pavlovian fear conditioning. Behav. Neurosci. 126, 404–422 (2012).

88. Zhang, X., Kim, J. & Tonegawa, S. Amygdala Reward Neurons Form and Store Fear Extinction Memory. Neuron 105, 1077–1093.e7 (2020).

89. Gerber, B. et al. Pain-relief learning in flies, rats, and man: basic research and applied perspectives. Learn. Mem. 21, 232–52 (2014).

90. Mayer, D., Kahl, E., Uzuneser, T. C. & Fendt, M. Role of the mesolimbic dopamine system in relief learning. Neuropsychopharmacology 43, 1651–1659 (2018).

91. Dinsmoor, J. a. Stimuli inevitably generated by behavior that avoids electric shock are inherently reinforcing. J. Exp. Anal. Behav. 75, 311–333 (2001).

92. Tremblay, L. & Schultz, W. Relative reward preference in primate orbitofrontal cortex. Nature 398, 704–708 (1999).

93. Winston, J. S., Vlaev, I., Seymour, B., Chater, N. & Dolan, R. J. Relative valuation of pain in human orbitofrontal cortex. J. Neurosci. 34, 14526–14535 (2014).

94. Riemensperger, T., Völler, T., Stock, P., Buchner, E. & Fiala, A. Punishment Prediction by Dopaminergic Neurons in Drosophila. Curr. Biol. 15, 1953–1960 (2005).

95. Felsenberg, J., Barnstedt, O., Cognigni, P., Lin, S. & Waddell, S. Re-evaluation of learned information in Drosophila. Nature 544, 240–244 (2017).

96. Jacob, P. F. et al. Prior experience conditionally inhibits the expression of new learning in Drosophila. Curr. Biol. 31, 3490–3503.e3 (2021).

97. Zhao, B. et al. Long-term memory is formed immediately without the need for protein synthesis-dependent consolidation in Drosophila. Nat. Commun. 10, (2019).

98. Bennett, J. E. M., Philippides, A. & Nowotny, T. Learning with reinforcement prediction errors in a model of the Drosophila mushroom body. Nat. Commun. 12, (2021).

99. Schindelin, J. et al. Fiji: An open-source platform for biological-image analysis. Nat. Methods 9, 676–682 (2012).

100. Kang, L. The image stabilizer plugin for ImageJ. (2008).

101. Placais, P.-Y. & Preat, T. To Favor Survival Under Food Shortage, the Brain Disables Costly Memory. Science (80-. ). 339, 440–442 (2013).

